# Distinct synovial fluid B-cell differentiation and activation profiles in rheumatoid arthritis

**DOI:** 10.64898/2026.09.24.754012

**Authors:** Wenqi Huang, Salim Ghannoum, Kittikorn Wangriatisak, Annelien Hooijsma, Maho Nakazawa, Lovisa Franzén, Stefan Petkov, Katerina Chatzidionysiou, Vivianne Malmström, Caroline Grönwall

## Abstract

CD11c+CD21- B cells are overrepresented in rheumatoid arthritis (RA). Here we delineate their functional state at site of inflammation and its molecular basis. Phenotypic profiling of paired blood and synovial fluid from patients with anti-citrullinated protein antibody (ACPA) seropositive RA (N=12), seronegative RA (N=10) and spondyloarthritis (N=8), demonstrated a striking enrichment of CD11c+CD21- B cells in synovial fluid in all three diseases (average of CD19+ cells: 67%, 49% and 51%). In ACPA-positive RA we identified increased proportions of CD11c+IgG+CD27^low^ cells and several stages of pre-plasmablasts. The CD11c+CD21- cells had a phenotype indicating BCR activation, proliferation, and antigen presentation. This B-cell profile was mirrored by a LAG3+CTLA4+TIGIT+ peripheral helper T-cell program in ACPA-positive RA, contrasting a CCR6+ Th17-skewing in seronegative RA. Single-cell spatial proteomics revealed a unique CD11c^high^CD21^low^ B-cell surface architecture including a BCR co-receptor-tetraspanin signature with CD72-CD81 co-localization, which was elevated in RA. CD11c+ cells also displayed CD44-CD52/CD59 and MHCII-CD40/CD84 immune-modules absent in other B-cell subsets.

In summary, CD11c+CD21- B cells are highly diverse, and while an activated antigen-experienced synovial profile was shared between disease groups, ACPA+ RA showed evidence of increased extrafollicular plasmablast differentiation. Surface receptor organization implicated a favoured BCR responsiveness, complement resistance and antigen presentation.

## INTRODUCTION

Rheumatoid arthritis (RA) is chronic inflammatory disease which is defined by synovitis and arthralgia. A large proportion of RA are classified to have seropositive disease which is associated with an autoimmune response and anti-citrullinated protein autoantibodies (ACPA) as well as anti-IgG Fc (RF, rheumatoid factor)[1]. A pivotal role of the adaptive immune system, B cells and T cells, in the pathogenesis of seropositive RA is solidified by certain HLA-DRB1 alleles being the strongest genetic risk factor for RA, directly implicating MHCII presentation and CD4+ T cells as essential [2]. Moreover, the clinical efficacy of both B cell depletion [3] and T cell co-stimulation blockade [4] further frames seropositive RA as driven by pathogenic T-B cell collaboration. For seropositive RA, several B cell repertoire and phenotypic shifts have been observed in the circulation, including an increase of CD27-negative cells and low mutated IgG, changes in Fab-glycosylation sites, and elevation of activation markers across different B cell subsets [5–9]. Studies have also reported aberrant BCR signalling with elevated phosphorylation of Syk (p-Syk) downstream of BCR in certain B cell populations [10–12]. CD72 is a regulator of BCR signalling that arise emerging interest in the context of BCR-signalling changes in autoimmunity. In SLE, selective loss of CD72 on DN2 B cells has recently been linked to enhanced p-Syk/p-Erk responses[13], but its role in RA has not been addressed.

One B cell phenotype that has received attention in multiple autoimmune and chronic inflammatory conditions, including in ACPA+ RA, is CD11c+ CD21- B cells. These cells can include both activated naïve (aNAV) if they are IgD+ CD27-, and atypical memory B cells or double-negative 2 (DN2) cells if they lack both IgD and CD27 [14–18]. CD11c+ CD21- cells, also denoted age-associated or autoimmune-associated B cells (ABCs) if positive for the transcription factor T-bet, have been reported to be expanded in seropositive RA blood and correlate with both joint destruction and disease activity [19]. Single cell transcriptomics studies have suggested that RA synovium CD11c+ DN2 cells are precursors to antibody secreting cells [17], while reports analysing peripheral blood and synovium ABCs show transcript profiles consistent with antigen presenting capacity [19].

T cell - B cell interaction is key in seropositive RA, with CD4+ T cells providing B cell help to generate ACPA responses with immunoglobulins carrying high levels of somatic hypermutations, and B cells presenting antigen to CD4+ T cells and thereby providing a forward loop to chronic autoimmune responses. Such interactions are less defined for seronegative RA and other forms of arthritis.

In the current study, we investigate blood and synovial B cell and T cell compartments across ACPA+ RA, ACPA-RA and spondyloarthritis (SpA), combining multiparameter flow cytometry, and single cell spatial proteomics. We find that CD11c+ CD21- B cells dominate the synovial B cell pool but that there are distinct differences in the profiles of both B cells and T cells between the groups. In ACPA+ RA, the CD11c+ CD27- synovial B cell profile was paired with a Tph response while ACPA-RA had synovial T cells skewed towards a CCR6+ Th17 phenotype. The CD11c+ CD21- cells in ACPA+ RA were further found to carry a differential surface protein spatial architecture that further distinguishes RA from healthy controls.

## MATERIALS AND METHODS

### Patients and samples

All RA patients fulfilled the American College of Rheumatology (ACR) / The European Alliance of Associations for Rheumatology (EULAR) classification criteria[20]. Synovial fluid (SF) samples were obtained from patients with ACPA-RA (n=10), ACPA+ RA (n=12), and spondyloarthritis (SpA, n=8) that were undergoing large joint arthrocentesis as part of clinical care at the Rheumatology Clinic, Karolinska University Hospital, Stockholm (Supplementary Table 1). Patient receiving B cell targeted treatment (e.g. Rituximab) <6 months before sample draw were excluded. Paired blood samples were available from 10 ACPA-RA patients, 9 ACPA+ RA patients, and 7 SpA patients. Seven of nine ACPA+ RA samples were RF+ while all ACPA-RA samples were RF-. In addition, peripheral blood samples from three ACPA+ untreated RA patients at disease onset and age-matched healthy blood donors (n=2) were included for single-cell spatial proteomics (Supplementary Table 1).

Peripheral blood mononuclear cells (PBMCs) and SF mononuclear cells (SFMCs) were isolated by Ficoll Paque density-gradient centrifugation (Cytiva) and cryopreserved until use. This study was approved by regional ethics committee and all participants provided informed consent in accordance with the Declaration of Helsinki.

### Flow cytometry

Samples from ACPA+ SF, ACPA-SF, SpA SF, and peripheral blood were always processed in parallel within the same experiment. Based on cell recovery, one to three flow cytometry panels were applied (Supplementary Table 2). For B cell phenotyping 5 × 10^6^ PBMCs or SFMCs were stained with fixable viability dye eFluor 506 (1:1000; Invitrogen) for 10 min, followed by staining with fluorochrome-conjugated antibodies (Supplementary Table 2). The gating strategies are shown in Fig. S1.

For T cell phenotyping 1 x 10^6^ of PBMCs or SFMCs were used for intracellular staining, cells were fixed and permeabilized using the FOXP3 Permeabilization Buffer Kit (eBioscience) according to the manufacturer’s instructions. Data were acquired on a 5-laser Cytek Aurora spectral flow cytometer and analyzed using FlowJo software (version 10.10.1) and R studio (4.5.2). The detailed description UMAPs generation in R is available in the Supporting Information. The gating strategies are shown in Fig. S2.

### Intracellular phosphoflow staining

PBMCs (3x10^6^ cells) were incubated with fixable viability dye eFluor 506 (1:1000; Invitrogen) for 10 min, washed twice with PBS, and surface-stained with antibodies cocktail (Supplementary Table 2) for 20 min. Cells were then washed and rested for 1 h at 37°C in RPMI supplemented with 2% FBS, 2 mM L-glutamine, 100 U/mL penicillin, 100 μg/mL streptomycin, and 10 mM HEPES.

Cells were subsequently stimulated with 20 μg/mL F(ab’)2 anti-IgM and anti-IgG (Jackson ImmunoResearch) and incubated at 37°C for 2 min or 5 min to measure p-Syk and p-Erk, respectively. For phospho-protein detection, cells were fixed, permeabilized, and stained with monoclonal antibodies specific for phosphorylated SYK (pY348) or ERK (pY204). Samples were acquired on a 5-laser Cytek Aurora spectral flow cytometer. The gating strategies are shown in Fig. S3.

### Proximity Network Assay

Proximity network analysis (PNA) was performed using the Pixelgen Proxiome Kit[21], Immuno 155 (PROXIMM001: as a pilot version which did not include CD79a), according to the manufacturer’s instructions. B cells were purified from PBMCs by negative selection using the B cell isolation Kit II (Miltenyi Biotec) and analyzed for three RA patients and two healthy controls. For RA paired total PBMC samples were also included. Proximity libraries were prepared according to the Pixelgen instructions. Briefly, 0.5-1 × 10^6^ cells were fixed with methanol-free paraformaldehyde (1% PFA fixation), followed by 0,5% BSA blocking, and subsequent incubation with the barcoded antibody panel in two steps. Protein proximity networks were generated by sequential padlock probe hybridization, padlock circularization, rolling circle amplification, and proximity ligation. After manual counting, 200-1,000 cells per sample were used for pre-amplification PCR and sample-index PCR, followed by SPRI bead purification (Beckman-Coulter Life science. Final libraries were quality controlled and quantified for the expected fragment size (∼277 bp) using Qubit dsDNA high sensitivity assay kit (ThermoFisher)) and gel electrophoresis, pooled equimolarly, and sequenced at the National Genomics Infrastructure (SciLifeLab, Stockholm, Sweden) on an Illumina NovaSeq X Plus. The eight libraries were pooled equimolarly and sequenced as a single pool on a 25B flow cell using the 300-cycle kit and the standard paired-end read setup, with dual 8 bp indices (i7 and i5) and a 15% PhiX spike-in, at a target sequencing depth of 250,000 read pairs per cell (approximately 2 × 10^9 read pairs in total).

### PNA data processing

Raw PNA sequencing data were processed with the nf-core/pixelator (Nextflow) pipeline. Downstream analyses used the pixelatorR package (v0.13.0) with Seurat (v5.2.1); plots were generated with ggplot2 (v3.5.1) and ComplexHeatmap (v2.22.0). Cells were retained if they had a normal Tau dispersion score, ≥20,000 UMIs, isotype control fraction <0.1%, and a dangling-node (k-core 1) fraction <0.5. Protein abundances were CLR-normalized, and UMAP embedding with Louvain clustering (resolution = 0.8) was applied to the first 18 Harmony-corrected PCs, yielding 8 clusters. For proximity analyses, per-cell background was removed with FilterProximityScores (background_threshold_pct = 0.00071; min_cells_count = 5); summarized analyses further required UMI >100 per marker and join count >5, with per-population mean log₂ ratio. Proximity-based UMAP and Louvain clustering used dims = 1:8 (resolution = 0.2). Details are provided in the Supporting Information.

### Statistics

Flow cytometry results were analyzed with the Prism 9 software (GraphPad) or R studio (4.5.2). Medians were compared by Mann-Whitney U test (two groups); Wilcoxon matched-pairs signed rank test (paired two groups) or Kruskal-Wallis (> 2 groups) with correction for Dunn’s multiple comparisons. Spearman’s test was applied to investigate correlations. P-values less than 0.05 were considered statistically significant. The performed statistical test for each analysis is indicated in the figure legends.

## RESULTS

### Synovial B cells have an activated phenotype in different arthritis subsets

To better understanding of B cells at site of inflammation we performed phenotypic analyses in paired peripheral blood (PB) and synovial fluid (SF) samples from patients with ACPA-positive rheumatoid arthritis (ACPA+ RA, PB=9; SF=12), ACPA-negative RA (ACPA-RA, PB=10; SF=10), and spondyloarthritis (SpA, PB=7; SF=8) with high-dimensional spectral flow cytometry. All ACPA-patients were also RF IgM negative and a majority of ACPA+ patients were RF positive (7 of 9). All patients had active disease that required a synovial fluid aspiration. Twelve patients were untreated at time of sampling (N=7 ACPA+ RA, N=1 ACPA-RA and N=4 SpA), five were receiving only MTX (N=1 ACPA+ RA, N=3 ACPA-RA and N=1 SpA) and eight were receiving TNFi with/without MTX (N=3 ACPA+ RA, N=4 ACPA-RA and N=2 SpA). Detailed information is shown in Table S1.

As expected, total B cell frequencies were lower in synovial fluid relative to paired blood across all three groups. Interestingly, substantial numbers of synovial B cells were detected also in the SpA group, and while synovial fluid from ACPA+ patients had a significantly higher B cell frequency than ACPA-patients, no statistical difference was seen compared to SpA patients (ACPA+ SF: 1,52%; ACPA-SF:0,68%; SpA SF:1,3%; Fig. 1A).

**Figure 1.**
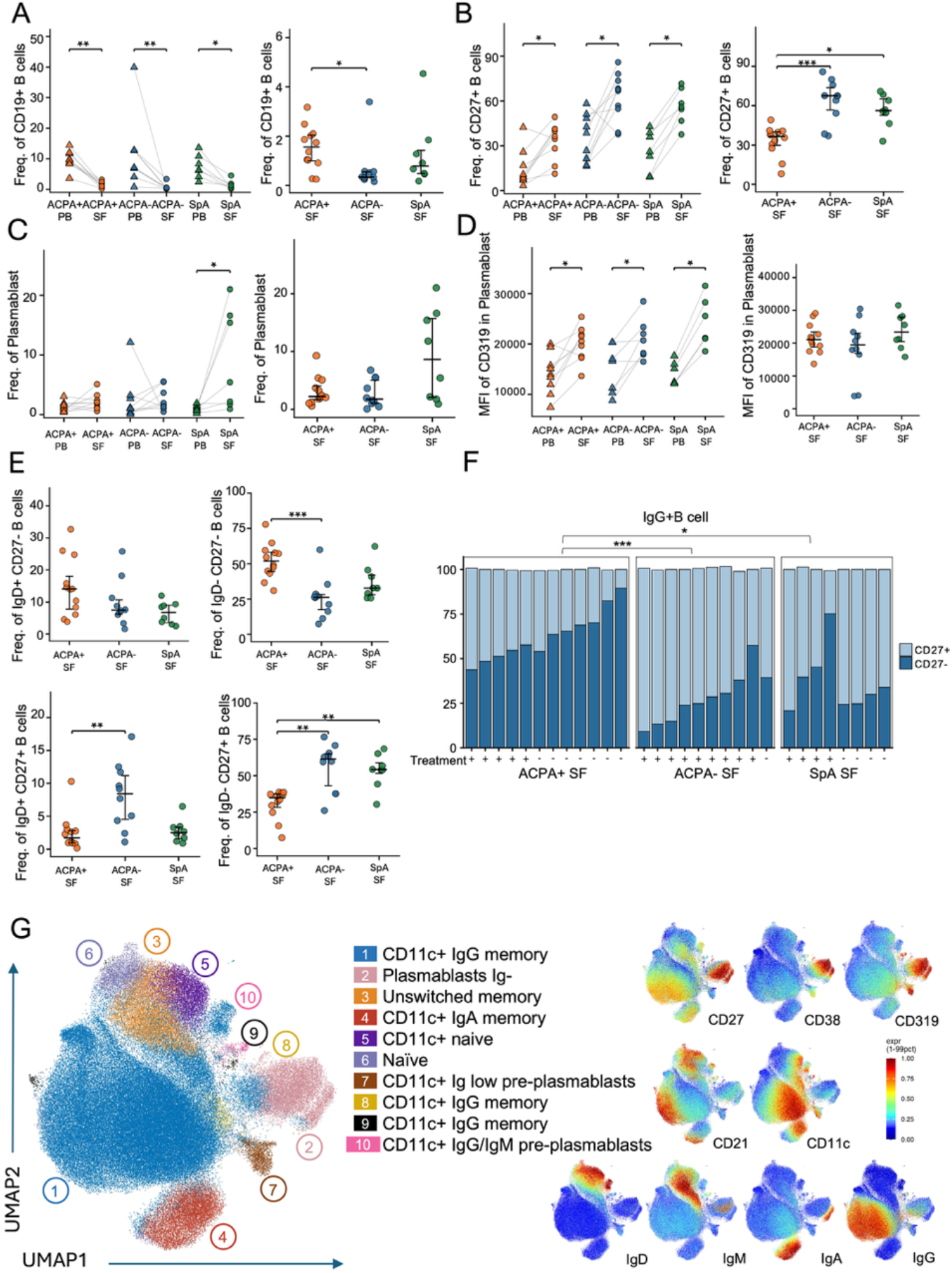
Synovial fluid B cells have reduced CD27 expression in ACPA-positive RA. B-cells were analyzed with spectral flow cytometry in paired peripheral blood (PB) and synovial fluid (SF) from ACPA+ RA (PB N=9, SF N=12), ACPA-RA (PB N=10, SF N=10) and SpA patients (PB N=7, SF N=8). Frequencies are shown of all CD19+ B cells (**A**), CD27+ B cells (**B**), CD27^high^ CD38^high^ plasma cells (**C**), Average MFI of CD319 on plasmablasts in paired PB and SF (left) and SF plasmablast between groups (right) (**D**) and IgD/CD27-defined subsets [naïve: IgD+ CD27-, unswitched memory IgD+ CD27+, switched memory: IgD-CD27+, and double negative: IgD-CD27-] (**E**). The left panels in A-D show paired PB and SF frequencies and the right panels compare proportions between disease groups in SF (right). **F**. Stacked bar graphs show the proportion of CD27+ and CD27-cells within IgG+ SF B cells for individual patients by disease group and annotated by treatment status (+, treated; -, untreated). **G**. Cluster analysis and UMAP visualization of SF B cells identified 10 distinct B cell clusters (left). Normalized MFI of lineage markers in the UMPA clusters are shown in the right panel with red indicating high expression. A summary of the observed cluster features is presented in Table 1. In the scatter plots, colors indicate: ACPA+ (orange), ACPA- (blue), SpA (green); PB, triangles; SF, circles. Paired samples are connected by grey lines. Horizontal bars indicate median ± IQR. Kruskal-Wallis test followed by Dunn’s multiple-comparisons was applied for inter-group SF comparisons; Wilcoxon matched-pairs signed-rank test was used for paired PB-SF comparisons. *P < 0.05, **P < 0.01, ***P < 0.001.

**Table 1.** Identified synovial B cell clusters.

| No | Cluster identity | CD11c | CD21 | CD27 | IgD | IgM | IgG | IgA | Other significant markers | Features |
| --- | --- | --- | --- | --- | --- | --- | --- | --- | --- | --- |
| 1 | CD11c+ IgG memory | +++ | - | ++ | - |  | +++ |  |  | Gradient of<br>higher CD11c<br>lower CD27<br>low surface Ig |
| 2 | Plasmablasts Ig- | - | - | ++++ | - |  | - | + | CD319, CD38+++ |  |
| 3 | Unswitched memory | - | ++ | ++ | +++ | +++ | - |  |  |  |
| 4 | CD11c+ IgA memory | +++ | - | +++ | - |  | - | +++ |  | Gradient of<br>higher CD11c<br>lower CD27<br>(no change in surface Ig) |
| 5 | CD11c+ naive | +++ |  |  |  |  |  |  |  |  |
| 6 | Naive | - | +++ | - | +++ | ++ | - |  | CD21, CD72 |  |
| 7 | CD11c+ Ig low pre-plasmablasts | +++ | - | + | - |  | - |  | CD319, CD38++ CD197, |  |
| 8 | CD11c+ IgG memory | +++ | - | + | - |  | ++ |  | CD43 |  |
| 9 | CD11c+ IgG memory | ++ | + | ++ | - |  | ++ |  | CD38+, CD10 |  |
| 10 | CD11c+ IgG/IgM pre-plasmablasts | +++ | ++ | ++ | - | ++ | ++ |  | CD319, CD38+, CD1c, CD185 |  |

In general, synovial fluid B cells showed higher expression and frequency of proliferation and activation markers (HLA-DR, CD95, CD71, CD86) than blood B cells in terms of MFI in total CD19+ B cells (Fig. S4a, b). Concurrently, synovial fluid also displayed lower expression and frequency of inhibitory and regulatory surface receptors, including CD72, CD73 and BTLA (Fig. S4a, b). We could however see higher expression and frequency of inhibitory marker TIGIT in the synovial CD19 B cell compared to peripheral blood. Moreover, SF B cells showed markedly reduced expression and frequency of the canonical homing receptors, CXCR5 and CCR7 (CD197), while CXCR3 was upregulated (Fig. S4a, b).

Compared to paired blood, synovial fluid B cells across all three groups showed a higher proportion of CD27+ cells and increased IgG and IgA expression, reflecting a shift toward memory B cells and plasmablasts in the joint (Fig. 1B left, Fig. S4a, b). Although memory B cells were the dominant population, plasmablast was seen in all subgroups but was most prominent in SF from SpA with 8.6% plasmablast compared to 1-2% in ACPA+ and ACPA-RA. (Fig. 1C). The SF CD19+ plasmablasts shared a common phenotypic signature in all three groups, with elevated CD38 and CD27 in combination with CD319 (SLAMF7), a marker of plasma cell commitment (Fig. 1C, D), paralleled by reduced BTLA and CD43 expression (Fig. S5A, B). Consequently, the synovial B cells had a higher activation, differentiation and proliferation state, and display evidence of migration.

### Synovial B cells in ACPA-positive RA exhibit loss of CD27 expression and expansion of class-switched CD27-negative populations

When further analyzing the overall composition of the non-plasmablast synovial B cells, we observed that on only 36% of B cells in ACPA+ RA patients displayed CD27 expression compared to 67% and 56% from ACPA-RA and SpA patients (Fig. 1B right). This CD27-expansion was predominantly explained by an elevated frequency of CD27- IgD-double-negative) B cells (52%) (Fig. 1E) and not from an increase in IgD+ CD27- B cells (14%). Correspondingly, the synovial B cells from ACPA+ patients also contained proportionally lower frequencies of both IgD-CD27+ switched memory (SM, 35%) and IgD+ CD27+ unswitched memory (USM, 1.5%) B cells compared with ACPA- (SM: 61%; USM:8%). (Fig. 1E).

Moreover, the synovial DN compartment in ACPA+ RA was enriched for class-switched cells as demonstrated by significantly higher abundance of both IgG+ and IgA+ among CD27- B cells compared with ACPA-RA and SpA SF B cells (Fig. 1F, Fig. S6). Taken together, these findings indicate that ACPA+ RA is characterized by a synovial B cell landscape dominated by CD27 low, class-switched populations - a pattern not observed in ACPA-RA or SpA.

### ACPA-positive synovial fluid is enriched for CD11c+ CD21- B cells with a predominant IgG+ CD27- phenotype and expansion of pre-plasmablasts

Unsupervised cluster analysis and dimensionality reduction by UMAP of the flow cytometry B cell data from all three patient groups further highlighted the phenotypic differences between the synovial fluid and blood compartment (Fig. 1G, 2A, Fig. S7, S8). Analysis of the synovial B cells revealed 10 different B cell clusters (Fig 1G) which could be compared to 15 clusters in a combined PB and SF analysis (Fig S7). All clusters were represented across different donors but with some individual differences (Fig S8A). The anatomical sites showed several distinct features where SF had a striking enrichment for CD11c+ clusters (Fig 1G, Fig S8A, Table 1). Low CD27 was detected as a gradient in CD11c+ IgG (cluster 1) and CD11c+ IgA+ clusters (cluster 4) as well as in several other atypical CD11c+ memory and pre-plasmablast clusters (cluster 5-9).

**Figure 2.**
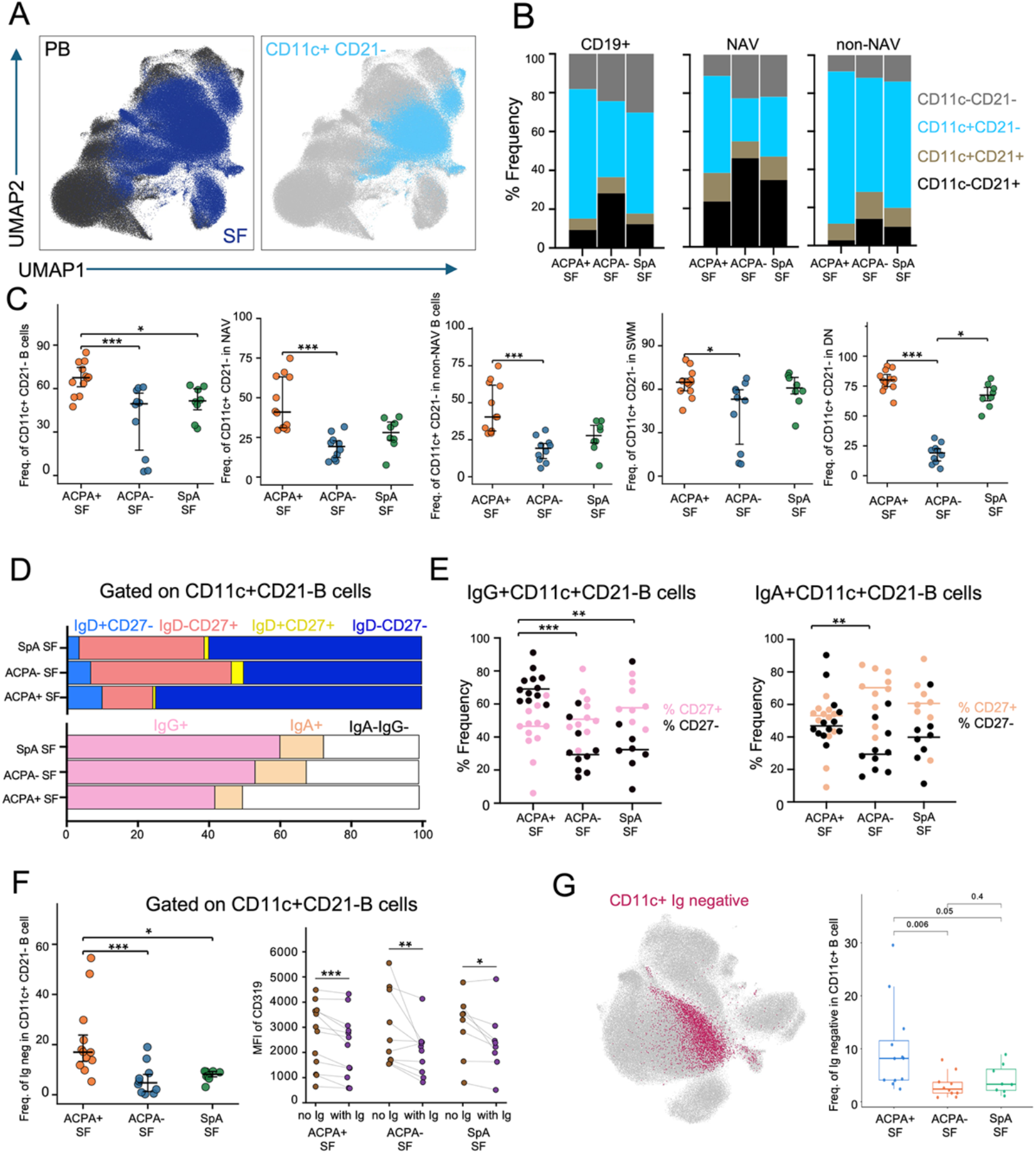
CD11c+CD21- B cells are overrepresented in synovial fluid with expansion of immature CD11c+ plasmablasts in ACPA+ RA. **A**. UMAP visualization of joint clustering of CD19+ B cells originating from peripheral blood (PB, black) or synovial fluid (SF, blue) and with the CD11c+ CD21- populations highlighted (right, light blue). **B**. Stacked bar graphs showing the proportional distribution of the four CD11c/CD21-defined subsets (CD11c- CD21-, CD11c+ CD21-, CD11c+ CD21+, CD11c- CD21+) within the CD19+ naïve (NAV) or non-naïve (non-NAV) SF compartments in three disease groups (ACPA+ RA, ACPA- RA and SpA). **C**. Frequencies of CD11c+ CD21- cells within total B cells and within individual subsets (NAV, non-NAV, switched memory [SWM] and double-negative [DN]) in SF between disease groups. **D**. Stacked bar graphs showing the proportion of IgD/CD27-defined subpopulations [IgD+ CD27-, IgD-CD27+, IgD+ CD27+, IgD-CD27-] (top panel) and immunoglobulin isotype surface positivity (IgG+, IgA+ or IgA-IgG-) in SF CD11c+ CD21- B cells in the three disease groups. **E**. Proportion of CD27+ versus CD27- cells within IgG+ (left) and IgA+ (right) CD11c+CD21- SF B cells in disease groups. **F**. Frequency of Ig-cells (IgA-IgG-IgM-IgD-) within CD11c+ CD21- B cells across SF disease groups (left) and the MFI of CD319 on Ig negative versus Ig positive in CD11c+ CD21- (right). Colors on the scatter plot indicate: ACPA+ SF (orange), ACPA- SF (blue), SpA SF (green). Horizontal bars indicate median±IQR. Kruskal-Wallis test with Dunn’s correction of multiple comparisons was applied for inter-group comparisons; Wilcoxon signed-rank test was used for paired comparisons. *P < 0.05, **P < 0.01, ***P < 0.001.

Manual gating confirmed that the CD11c+ CD21- B cell population constituted the predominant synovial B cell subset in all three groups (Fig. 2B, bar graphs average: ACPA+: 67%; ACPA-: 49.6%; SpA: 51%). Still, ACPA+ patients showed a significantly higher frequency of synovial CD11c+ CD21- B cells compared with both ACPA- and SpA patients and across multiple B cell subsets, including both naïve and non-naïve populations (Fig. 2B, C).

Within the CD11c+ CD21- compartment, IgG expressing CD27- B cells were especially more prominent in SF of ACPA+ RA (Fig 2D, E). Also, the IgA+ CD11c+ CD21- B cells in ACPA+ RA contained a significant proportion of CD27-, but the difference was not as striking as for IgG. Such class-switched CD11c+ CD21- B cells were less pronounced in the other disease groups. In addition, the subset of CD11c+ CD21- cells without detectable surface immunoglobulin was higher observed in ACPA+ SF (Fig 2F). Those surface immunoglobulin-negative B cells further had higher expression of CD319 (SLAMF7) and CD197 (CCR7) and lower expression of CD45RB compared with Ig+ CD11c+ CD21- B cells (Fig 2F, Fig S9B). Hence, we postulate that these cells represent CD11c+ CD21- cells with pre-plasmablast features. When comparing with the UMAP analysis, we could observe a gradient loss of Ig within the largest CD11c+ CD21- IgG+ cluster (cluster 1) (Fig 2G) which coincided with loss of CD27 expression. In addition, there were low surface Ig expression in a distinct CD11c+ CD21- CD38++ CD319+ CD27^low^ CD197+ cluster of immature pre-plasmablasts (cluster 7). Notably, no correlation was seen between expansions of immature CD11c+ pre-plasmablasts and the regular CD11c- mature plasmablast cluster (cluster 2).

In summary, ACPA+ RA was characterized by an increased synovial CD11c+ IgG+ CD27^low^ compartment which was associated with a gradient gain of a pre-plasmablast phenotype characterized by loss of surface immunoglobulin and increase of CD319.

### TNFi treatment associates with lower synovial proportions of CD11c+ B cell

In all patient groups we included individuals with and without ongoing MTX and TNFi treatment. When comparing treatment groups we found several significant differences. Treated RA patients displayed lower frequencies of double negative (IgD- CD27-) SF B cells compared with untreated patients (29% vs 56%, respectively) and correspondingly higher frequencies of IgD- CD27+ switched and IgD+ CD27+ unswitched memory populations (Fig. S10A). In SF from untreated RA patients, IgG+ and IgA+ B cells were more often CD27- than CD27+, whereas treated patients showed the reverse pattern, with IgG+ and IgA+ cells predominantly falling within the CD27+ memory fraction (Fig. S10B, C). Within the CD11c+ CD21- compartment, treatment with MTX/TNFi was associated with a lower frequency of these cells in RA SF (median 50% vs 68% in untreated), a difference that was preserved when the analysis was restricted to the naïve, DN and SWM compartments (CD11c+ CD21- among naive B cells: 23% vs 56%; among DN B cells: 61% vs 81%; among SWM B cells: 58% vs 65%; Fig. S10D). The blood compartment displayed the opposite trend: treated patients had a higher frequency of CD11c+ CD21- cells within the naive compartment (Fig. S10E). There’s no difference between untreated and treated groups in SpA (Fig. S10F).

### Synovial CD11c+ CD21- B cells display a lowered BCR activation threshold and an activated surface phenotype, most pronounced in ACPA+ RA

Given the predominance of CD11c+ CD21- B cells in SF, we further examined their functional phenotype relative to the rest of the B cell pool (Fig. 3A-C, Fig. S11). The CD11c+ CD21- subset had an enrichment of cells with elevated expression of CD19, CD20, IgG, IgA, CD43, CD71, CD95, and TIGIT together with reduced IgD, IgM and CD24 which was observed in all the groups (Fig. 3C, Fig, S11).

**Figure 3.**
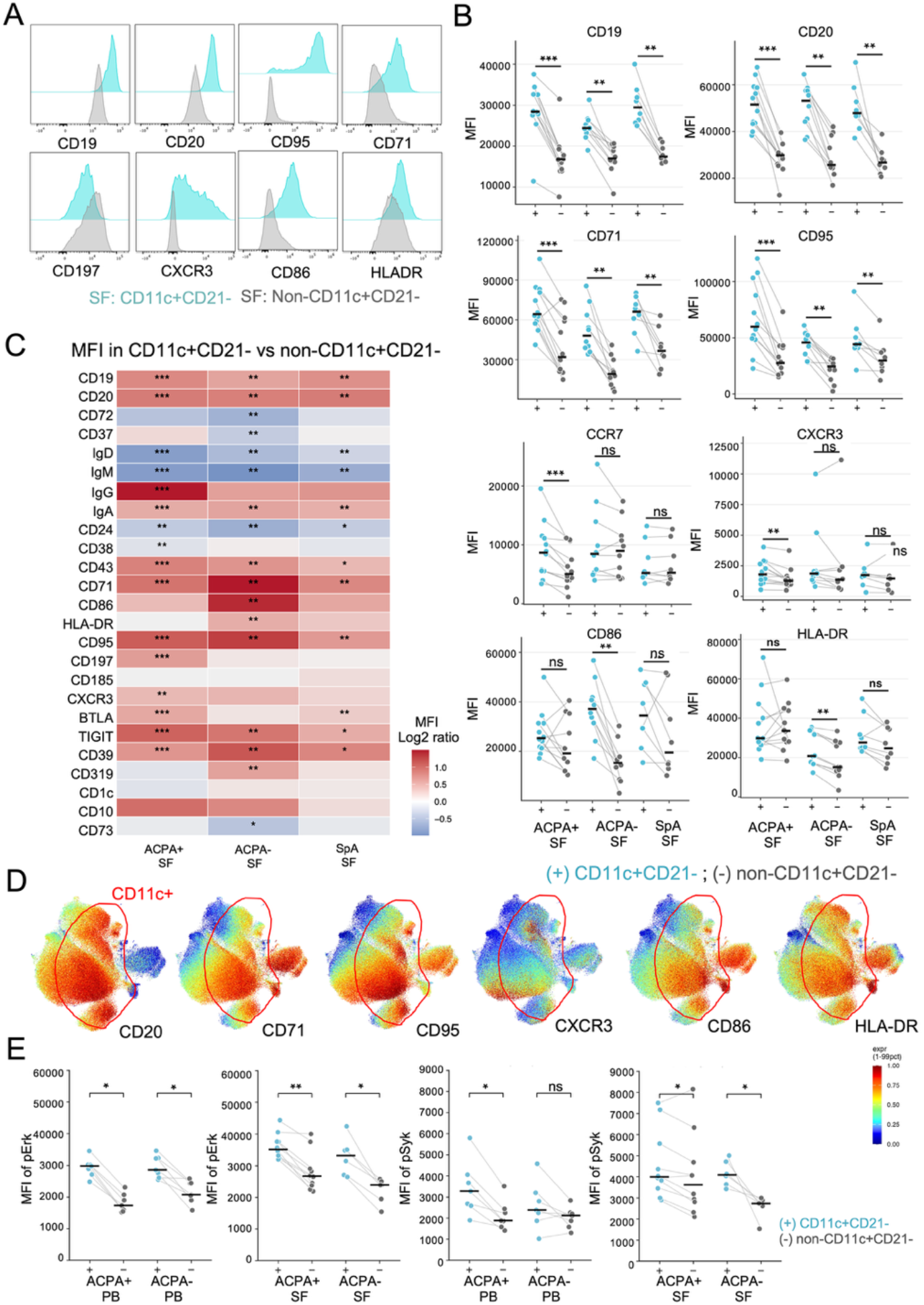
CD11c+CD21- B cells display a distinct activated phenotype and heightened BCR-proximal signaling relative to non-CD11c+CD21- B cells. **A.** Representative flow cytometry density histograms overlaying SF CD11c+CD21- (cyan) and non-CD11c+CD21- (grey) B cells for MFI of indicated markers. **B.** MFI of CD19, CD20, CD71, CD95, CCR7, CXCR3, CD86 and HLA-DR in CD11c+CD21- (+) versus non-CD11c+CD21- (-) B cells within each synovial fluid (SF) disease group; paired values are connected by grey lines. **C.** Heatmap of the log₂ MFI ratio (CD11c+ CD21- versus non-CD11c+ CD21-) for the indicated surface markers in each SF disease group. Red indicates higher expression in CD11c+ CD21- cell and blue lower expression compared to non-CD11c+ CD21-. **D.** Distribution of normalized MFI of activation markers in different B cell populations visualized in UMPA analysis. Red indicates higher expression. **E.** Phosphoflow analysis after BCR stimulation of peripheral blood or synovial B cells. MFI of phospho-ERK (pERK) and phospho-SYK (pSYK) in CD11c+ CD21- (+) versus non-CD11c+ CD21- (-) B cells from ACPA+ and ACPA- patients. Wilcoxon signed-rank test was applied for paired CD11c+CD21- versus non-CD11c+ CD21- comparisons, with FDR correction where indicated. *P < 0.05, **P < 0.01, ***P < 0.001; ns, not significant.

Comparison of SF CD11c+ CD21- cells between ACPA+ and ACPA- RA revealed that those from ACPA+ RA exhibited higher expression of activation and survival markers including CD95, CD71, HLA-DR, CD19, CD39, and CD43 relative to SF B cells from ACPA- RA. Conversely, CD86 expression was instead higher in ACPA- SF (Fig, S12). Interestingly, CD72 was expressed at a higher level in CD11c+ CD21- cells in ACPA+ patients compared to ACPA- which was primarily originating from a difference in activated naive cells (Fig, S12).

UMAP cluster analysis demonstrated that higher CD71, CD95, CD86 and HLA-DR largely coincided with increasing CD11c expression (Fig 3D). It should be noted that also cells with a pre-plasmablast phenotype displayed high CD86 and HLA-DR. CXCR3 on the other hand showed mostly an inverse but distinct pattern.

The high activation profile of CD11c+ CD21- SF cells prompted us to further evaluate whether this population exhibited altered BCR activation threshold and increased downstream signalling. Flow cytometry to detect phosphorylated SYK (p-Syk) and phosphorylated ERK (p-Erk) was performed following in vitro anti-immunoglobulin stimulation of B cells isolated from paired PB and SF samples (Fig. 3E; Fig. S13A, B). In general, SF B cells exhibited higher frequencies of p-Erk+ and p-Syk+ cells compared with matched PB (Fig. S13A, B). Importantly, CD11c+ CD21- B cells exhibited higher p-Syk and p-Erk compared with non-CD11c+ CD21- cells across all patient groups (Fig. 3E), indicating intrinsically heightened BCR signaling potential of this subset. CD72^low^ cells displayed higher HLA-DR, CD86, CD95, and CD71 expression as well as elevated p-Syk and p-Erk induction relative to CD72+ cells (Fig. S14A-C). The signalling differences were most pronounced and consistent in the aNAV subset (Fig. S14C).

### Synovial T cell phenotypes differ between ACPA-positive RA, ACPA-negative RA and SpA, with divergent Tph and Th17 profiles

Given the B cell alteration between ACPA+ and ACPA- SF, we next examined whether T cell populations were concordantly altered in a manner consistent with differential T-B cell crosstalk. UMAP analysis shown T cells are different in between PB and SF (Fig. 4A). Within SF, ACPA+ patients surprisingly had a lower frequency of CD4+ T cells and, a higher frequency of CD8αβ+ T cells compared to ACPA- patients (Fig. 4B). Furthermore, higher GZMK+ CD8αβ+ T cells were found in ACPA+ SF than ACPA- SF (Fig. 4B). Manual gating analysis of paired PB and SF T cells showed that, in all three groups, SF had lower proportion of naïve T cells and were enriched for memory populations, Tph and Treg subsets (Fig. 4C and Fig. S15).

**Figure 4.**
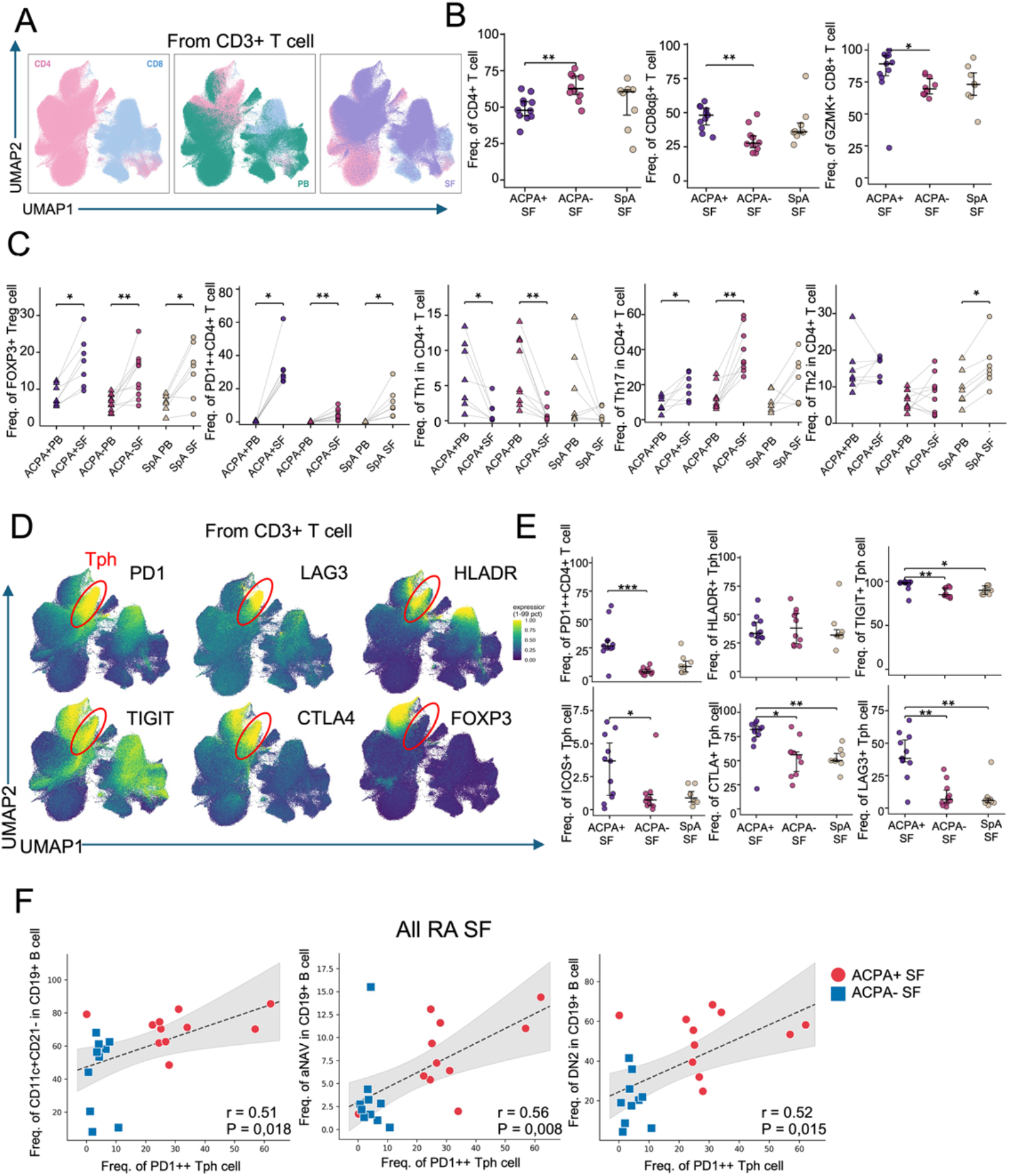
ACPA-positive synovial fluid is enriched for PD1++ Tph cells with a distinct checkpoint profile that correlates with the CD11c+CD21-, activated-naïve and DN2 B cell compartments. **A.** UMAP visualization of CD3+ T cells colored by CD4 versus CD8 lineage (left) and by origin (PB, SF). **B.** Frequency of CD4+ T cells, CD8αβ+ T cells and GZMK+CD8+ T cells in SF between disease groups (right). **C.** Paired PB-SF analysis of the proportion of Treg (FOXP3+), Tph (PD1++CD4+), Th17 (CXCR3-CCR4-CCR6+), Th1 (CCR6-CCR4-CXCR3+) and Th2 (CCR6-CXCR3-CCR4+) cells within CD4+ T cells, in all disease groups (ACPA+ RA, ACPA- RA and SpA). **D.** UMAP visualization of CD4+ T cells highlighting the PD1++ and Foxp3+ populations with MFI expression of HLA-DR, CTLA4, LAG3 and TIGIT. **E.** Frequencies of PD1++CD4+ T cells and of HLA-DR+, TIGIT+, ICOS+, CTLA4+ and LAG3+ cells within PD1++ Tph cells in SF between disease groups (right). **F.** Spearman correlation analysis between the frequency of PD1++ Tph cells and the frequency of all CD11c+ CD21- (r = 0.51, P = 0.018), CD11c+ naïve cells (activated-naïve, aNAV; R= 0.56, P = 0.008) and CD11c+ DN cells (DN2, R= 0.52, P= 0.015) cells within CD19+ B cells, including all RA SF samples; ACPA+ SF, red circles; ACPA- SF, blue squares; shaded bands indicate the 95% confidence interval. In the scatter plots the colors indicate: ACPA+ SF (purple), ACPA- SF (red), SpA SF (yellow); PB, triangles; SF, circles. Horizontal bars indicate median±IQR. Kruskal-Wallis test with Dunn’s correction for multiple-comparisons was applied for inter-group SF comparisons; Wilcoxon matched-pairs signed-rank test was used for paired PB-SF comparisons; Spearman rank correlation was used in (D). *P < 0.05, **P < 0.01, ***P < 0.001.

Th1 frequencies were consistently lower in SF than in paired PB. In contrast, the PB-to-SF enrichment of Th17 cells was most pronounced in ACPA- patients, whereas enrichment of Th2 cells was observed exclusively in SpA SF (Fig. 4C), suggesting a disease-specific pattern of synovial T cell accumulation. Interestingly, we found the frequency and expression of CXCR3 increased in SF T cell compared with blood while it was decreased in SF B cell compared those from blood (Fig. S16).

Synovial Tph cells were enriched in ACPA+ RA patients compared with ACPA- and SpA (Fig. 4E). These Tph cells expressed higher levels of HLA-DR, CTLA4, ICOS, LAG3, and TIGIT (Fig. 4D, E). When data from ACPA+ and ACPA- SF were pooled, the frequency of PD1++ Tph cells correlated positively with CD11c+CD21- B cell and CD11c+ aNAV and DN2 B cells (Fig. 4F, Fig. S17).

### CD11c^high^CD21^low^ B cells show altered spatial receptor structures in proximity to CD72, CD44 and MHCII, distinguishing them from other B-cell subsets

To investigate cell-surface protein organization in B cells from patients with RA, we applied the Proximity Network Assay (PNA) on purified B cells obtained from three untreated ACPA+ RA patients and two healthy controls (HC) (Fig. 5A). With the PNA technology >150 different proteins were used, which by proximity ligation and next generation sequencing give a computational reconstruction of single cells and a representation of the surface interactive proteome. Extensive quality control analysis confirmed high-quality data (Fig. S18,19). With unsupervised clustering we subseted B cells into 8 clusters (Fig. 5B). The identification of B cell subsets was consistent to what we found by flow cytometry analysis of the same samples (Fig. S19). Two PNA clusters (cluster 4: naïve and cluster 6: mixed unswitched/ switched memory) had a CD199 (CCR9) and CX3CR1 signal, both of which are predominantly intracellular in B cells. Hence, we speculate that these clusters may be a result of permeabilization and staining bias. Still and intriguingly, a distinct PNA cluster of CD11c^high^ CD21^low^ cells (cluster 7) were found. All eight clusters were detected in every donor, with no cluster restricted to an individual sample or disease group, although their relative proportions varied (Fig. 5C, D).

**Figure 5.**
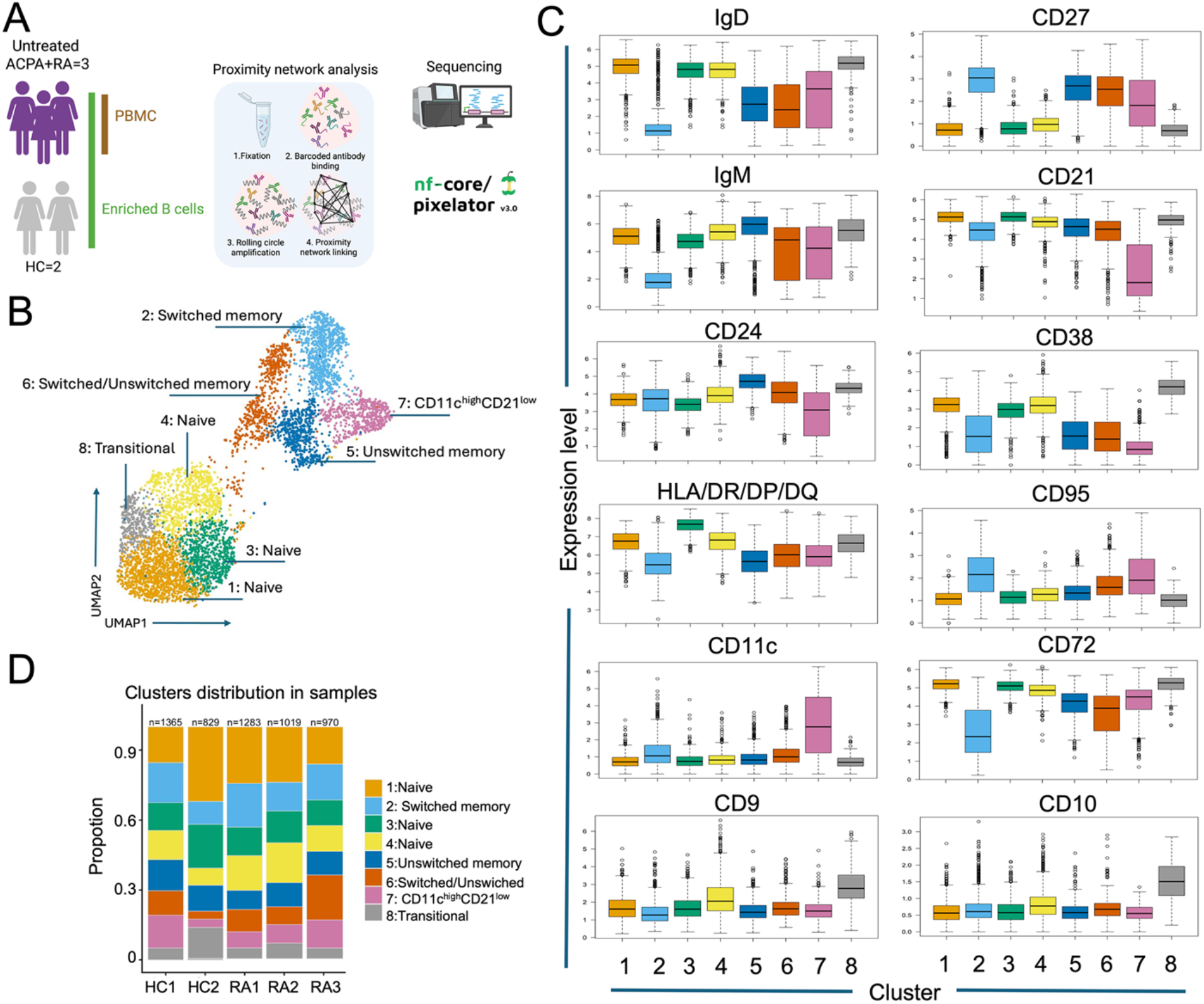
Proximity network analysis for single-cell cell-surface proteome profiling of RA B cells. **A.** Schematic of the experimental workflow. Peripheral blood mononuclear cells (PBMCs) and magnetically negative selection enriched B cells from three untreated ACPA-positive RA patients (RA1-RA3) and enriched B cells from two healthy controls (HC1, HC2) were processed using the Pixelgen Proxiome Cell-Surface Immuno-155 kit. The resulting libraries were sequenced on an Illumina platform, and reads were processed with nf-core/pixelator (v1.0). **B**. UMAP embedding of all B cells based on cell-surface protein abundance, colored by unsupervised cluster identity. Eight B cell clusters were resolved and annotated according to their canonical marker profiles: naïve (clusters 1, 3 and 4), switched memory (cluster 2), unswitched memory (cluster 5), switched/unswitched memory (cluster 6), CD11c^high^ CD21^low^ (cluster 7) and transitional (cluster 8). **C**. Box plots showing the expression of twelve canonical B-cell markers (IgD, CD27, IgM, CD21, CD24, CD38, HLA-DR/DP/DQ, CD95, CD11c, CD72, CD9 and CD10) across the eight clusters defined in (B). Box plots indicate the median (center line), the 25th-75th percentiles (box) and 1.5× the interquartile range (whiskers); individual cells beyond the whiskers are shown as dots. **D**. Stacked bar plot showing the relative proportion of each cluster per sample.

The resulting proximity matrix revealed a dense colocalization among components of the BCR signalling complex and associated tetraspanins. Cluster-resolved network representations of BCR co-receptors (CD72, CD19, CD81, CD82, CD21, CD37 and CD22) demonstrated that the BCR-tetraspanin module formed a stronger network in CD11c^high^ CD21^low^ cells (cluster 7) than in any other B-cell subset (Fig. 6A, B). Among these interactions, colocalization of the inhibitory co-receptor CD72 with the tetraspanin CD81 was the most pronounced feature of cluster 7. Per-sample mean log2 ratios for the CD72-CD81 pair were higher in CD11c^high^ CD21^low^ cells than in any other cluster (Fig. 6B, C). Importantly, those CD72-CD81 protein paired score was significantly higher in cells from RA cluster 7 cells relative to cells from healthy donor cluster 7 cells (P= 0.00051; Fig. 6D). Those findings were confirmed by manual gating analysis (Fig. S20).

**Figure 6.**
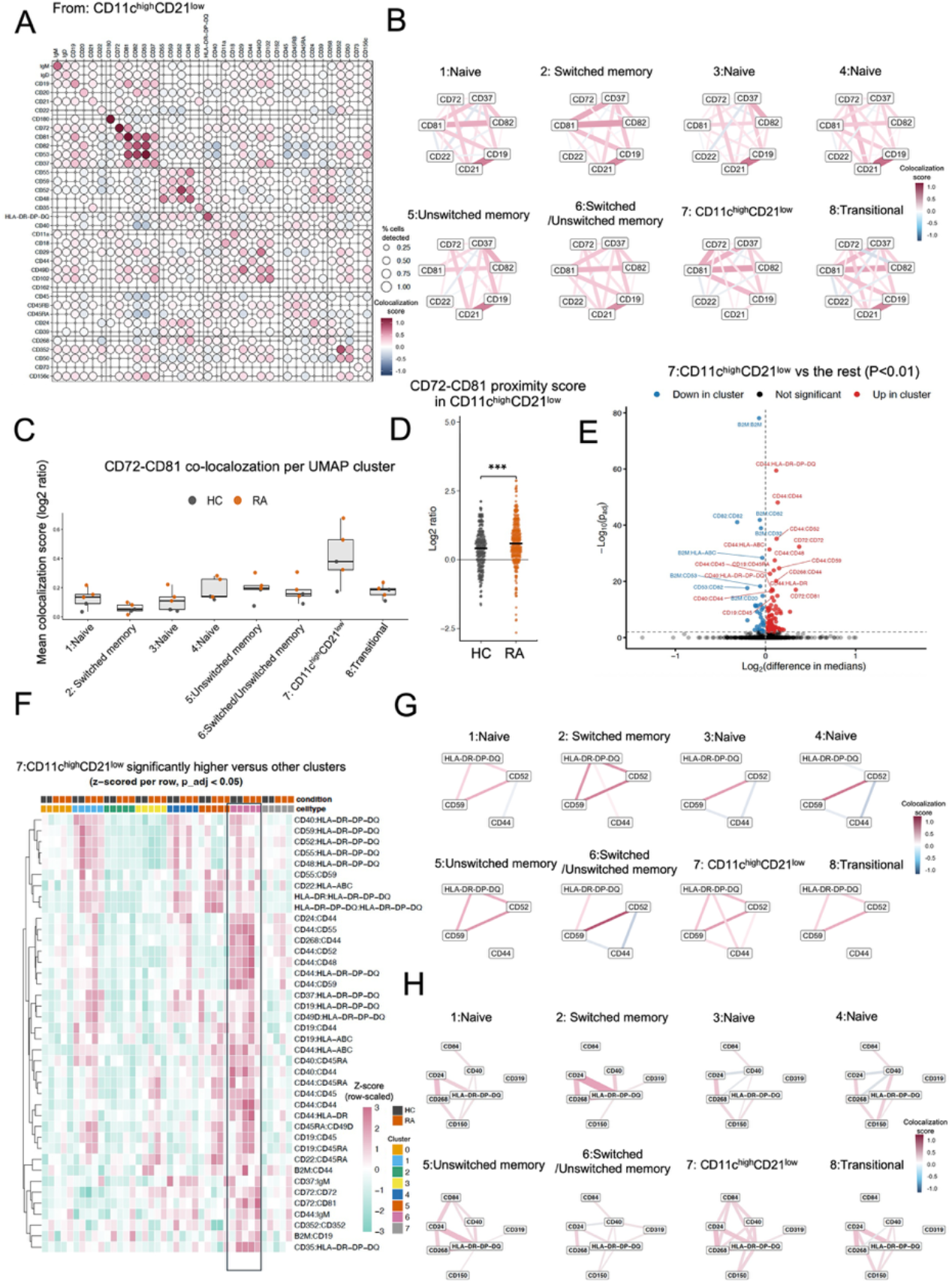
Proxiome profiling reveals unique receptor spatial architecture on CD11c+ B cells and alteration in ACPA+ RA compared to controls. **A.** Dot-plot heatmap of pairwise protein proximity scores across the eight B cell UMAP clusters, summarized as the mean log₂ ratio from PNA (left), dot size reflects the fraction of cells in which the pair was detected (pct_detected); **B.** Per-cluster network graphs of tetraspanin- and BCR-complex-associated markers (CD72, CD19, CD81, CD82, CD21, CD37, CD22; right). Width and edge color encode the mean log₂ ratio (red, colocalization; blue, segregation). Proximity data were pre-filtered for individual marker counts > 100 and join count > 5. **C.** Box plot of the mean CD72- CD81 colocalization score (log₂ ratio) per UMAP cluster (left; each point represents a per-sample mean; HC, grey; RA, orange), and per-cell CD72-CD81 proximity scores within the CD11c^high^ CD21^low^ cluster comparing all cells from healthy donors (HC) and RA patients (right; P = 5.1 × 10⁻⁴) shown in **D-E.** Volcano plot from Differential Proximity Analysis (DPA) comparing the CD11c^high^ CD21^low^ cluster against all remaining B cell clusters. The x-axis shows the difference in medians of the log₂ ratio and the y-axis -log₁₀ (adjusted P value). Pairs with significantly higher proximity in CD11c^high^ CD21^low^ cells are shown in red, those with lower proximity in blue; the dashed line indicates the significance threshold (P < 0.01). **F.** Heatmap of protein pairs with significantly higher proximity in the CD11c^high^ CD21^low^ cluster than in the remaining clusters (z-scored per row, adjusted P < 0.05), annotated by condition (HC, RA) and by cluster (1-8). **G.** CD44-centric proximity analysis in UMAP clusters: per-cluster network graphs of pairwise proximity among CD44, CD59, CD52 and HLA-DR/DP/DQ. **H.** MHC-II-centric proximity analysis across UMAP clusters: per-cluster network graphs of pairwise proximity among CD40, CD84, CD24, CD268, CD150, CD319 and HLA-DR/DP/DQ. (In G and H, edge width and colour encode the mean log₂ ratio, and proximity data were filtered as described in B.)

Differential proximity analysis of cluster 7 against all remaining B-cell clusters combined confirmed the CD72-CD81enrichment (P< 0.01) but also identified additional pairs dominated by CD44-centric interactions, including CD44-MHCII (detected with pan-HLA/DR/DP/DQ), CD44- CD52 and CD44-CD59 (Fig. 6E, F). Network analysis and paired analysis of CD44, CD52, CD59 and MHCII confirmed that the CD44-module has a configuration in cluster 7 that is different from all other B-cell subsets with higher colocalization of CD44-CD52, CD44-CD59, CD44-MHCII (Fig. 6G). CD44 mediates cell interactions and may by co-localization with CD52 and CD59 mediate changed cell-adhesion and complement resistance properties of CD11c+ cells.

Furthermore, a third module of significantly enriched pairs (P < 0.05; Fig. 6H) converged on the costimulatory molecule CD40 and on MHCII molecules. Network analysis of CD40, CD84, CD319 (SLAMF7), CD268 (BAFFR), CD150 (SLAMF1), CD24 and pan-MHCII revealed an MHC-centric pattern that was specific to cluster 7 and different from the other B-cell subsets. Especially CD319-MHCII, CD40-MHCII, CD84-MHCII, CD40-CD84 and CD40-CD268 co-localization was increased in CD11c^high^CD21^low^ B cells compared to the other cell clusters (Fig. 6H). A row-wise z-scored heatmap of the top significantly differential pairs (P< 0.05) recapitulated these findings, with CD44-, MHCII -containing pairs constituting the dominant proximity signature (Fig. 6F).

Consequently, the CD11c+ B cells displayed distinct surface receptor architecture which can be hypothesized to reflect increased BCR activity, antigen-presentation ability, complement resistance and tissue retention.

## DISCUSSION

In this investigation we provide a head-to-head comparison of the synovial fluid B cell compartment in seropositive and seronegative RA as well as in spondylarthritis. We find both interesting similarities and differences. A central finding was a dominance of CD11c+CD21- B cells in synovial fluid. ACPA+ RA showed a distinct synovial B cell phenotype with an augmented activation profile, a high proportion of IgG+ CD27- CD11c+CD21- cells and increased CD11c^high^ plasmablast-like cells with lost surface immunoglobulin. We provide further evidence that CD11c^high^ CD21^low^ cells have a distinct elevated activation state with unique spatial receptor organization supporting increased BCR activation, complement resistance, and antigen presentation. Altogether our data implicate that synovial B cells from ACPA+ RA have a strong trajectory from high antigen presentation in CD27^low^ IgG+ cells towards an immature plasmablast phenotype with no / low surface immunoglobulin.

Several distinct CD11c+ pre-plasmablast states were identified in ACPA+ RA. These CD11c+ pre-plasmablasts had acquired CD319 (SLAMF7), indicating entry into the plasma cell program [22]. Notably, in the SpA group, had CD11c- plasmablast expansions without many CD11c+ Ig low cells, suggesting a differential pathway in this disease group. Moreover, when comparing plasmablasts between blood and synovial fluid the regular CD11c- plasmablasts had higher expression of CD319 in all groups, indicating a more plasma cell differentiated state in the joint. This is consistent with RA synovial tissue harbouring ectopic lymphoid structures (ELS) [23] and ongoing plasma cell differentiation which has been detected already at diagnosis [24].

Another striking feature of the ACPA+ SF B cells was an expansion of B cells lacking CD27, with most of IgG+ cells residing in the CD27- “double-negative” gate. We have previously reported a similar B cell profile in circulating B cells in both ACPA+ RA and ACPA+ pre-clinical RA without arthritis [6, 25]. These cells have also been reported to have low somatic hypermutations compared to conventional CD27+ IgG+ cells in RA circulation [7, 26] and may be consistent with a bias towards extrafollicular immune responses [27]. Indeed, a majority of these IgG+ CD27- cells were CD11c+ CD21-. Indeed, we found that the synovial B cell compartment. The CD11c+ CD21- B cells are known to include both activated naïve (aNAV) and atypical memory/DN2 cell which are intensely studied in the context of infection and autoimmunity[28]. Circulating CD11c+ activated naïve B cells have been suggested to promote T cell responses to citrullinated antigen during the transition to clinical disease [29]. The nomenclature and functional properties of CD11c+ CD21- memory B cell is a topic of active discussion [27]. In our data, the phenotype of the overrepresented CD27- CD11c+ CD21- synovial cells intersected with atypical memory or DN2 cells although it should be noted that our panel did not include intracellular staining for T-bet [28]. In SLE, DN2 cells have been proposed to drive pathogenic autoantibody production[18]. In RA blood, frequencies of these cells have been found to be associated with more erosive disease[19] and to predict response to costimulatory blockade by CTLA4-Ig/abatacept [30]. Reduced frequencies of CD11c+ CD21- B cells following abatacept therapy has further been proposed to be linked to decreased Tph cells [31]. In our cohort, Tph cells were as expected especially elevated in ACPA+ SF and were characterized by markedly higher levels of the co-inhibitory receptors LAG-3, CTLA- 4 and TIGIT, a signature of chronic, antigen-driven activation[32]. This phenotype parallels the reported Tph-DN2 axis in transcriptomic data [17]. In line with this, Tph frequencies in our samples were associated with aNAV and DN2 frequencies.

Whether the CD11c+ CD21- B cells B cells act mainly as antibody precursors or also as antigen-presenting partners for T cells remains open: in RA synovial tissue, B cells with an atypical memory phenotype have been suggested to both represent progenitors of antibody-secreting plasmablasts as well as B cells with enhanced antigen-presenting capacity [17, 19]. Here we demonstrate that both features are indeed found in synovial fluid, but ACPA+ RA have a particularly strong trajectory towards antibody production. In comparison with published synovial tissue data, the CD11c+ compartment appears considerably more expanded in synovial fluid than in synovial tissue[17, 33], although differences in methodology mean that the data are not directly comparable. Moreover, high expansion of both Tph and CD11c+ pre-plasmablasts implicate a polyclonal response. Hence, this is unlikely to reflect a distinct antigen-specific response and may suggest more unspecific polyreactive features of the reaction.

ACPA- disease diverged on the T-cell side, with an enrichment of CCR6+ Tph17 and effector-memory CCR6+ CD4+ T cells in ACPA- RA pathogenically closer to a IL-23/IL-17 mechanism than to classical ACPA+ RA. Given that IL-17A blockade has shown only modest activity in unselected RA clinical trial[34], stratified studies will be required to test whether ACPA- subsets characterised by high CCR6+ Tph17 signatures might benefit preferentially.

The high CD11c expression in the synovium may also reflect migratory properties as the α-chain of the CD11c/CD18 (αXβ2, CR4) integrin mediates adhesion to inflamed vascular endothelium and transendothelial migration [35, 36]. A tight transcriptional concordance between peripheral-blood and synovial tissue T-bet+ CD11c+ B cells suggesting that the two compartments may reflect a common pool of trafficking cells[19]. However, BCR analyses of total B cells have reported limited shared clonotype lineages and expanded clones shared between compartments [37] [38]. Moreover, the direction and amplitude of trafficking between blood, synovial fluid and synovial tissue remains unclear. The potential association with synovial tissue pathotypes or cellular profiles merits investigation [40, 41]. In this context, the divergent CXCR3 pattern in the two lymphocyte compartments is of interest: relative to paired blood, synovial fluid B cells displayed higher CXCR3 whereas synovial T cells showed the opposite. As the interferon-induced ligands CXCL9- 11 are abundant in the rheumatoid joint[42], constitutively CXCR3-expressing T cells may lose surface receptor through ligand-driven internalisation after recruitment[43], whereas B cells, which express little CXCR3 at rest, appear to acquire it locally upon activation.

The hyperresponsive state of CD11c+ CD21- B cells have previously been documented in different RA ABCs [16, 19]. Our data investigate the molecular origin of this phenotype and suggest that it is associated with lower CD72 expression in non-naïve CD11c+ cells accompanied by altered CD72-BCR receptor spatial organisation. CD72 is classically regarded as an inhibitory BCR co-receptor[45]. CD72 is ITIM-bearing and recruits SHP-1 to dampen BCR signalling[46–48]. Engagement of its ligand CD100 (SEMA4D) dissociates the CD72-SHP-1 complex and relieves this inhibition, thereby promoting B cell activation[49–51]. Our data confirms increased BCR downstream pSyk and pErk in CD11c+ CD72^low^ cells compared to CD72^high^ cells, especially in activated naïve cells. In SLE DN2 cells, TLR7 stimulation selectively downregulates CD72[18], and reduced CD72 expression on DN2 B cells has recently been linked to disease activity and rituximab resistance[13].

BCR signal strength is governed by surface co-clustering-BCR micro-aggregation, CD19/CD21 recruitment, tetraspanin-organised microdomains and the spatial disposition of inhibitory receptors relative to the BCR signalosome[46, 52–54] the inhibitory function of CD72 depends not only on its abundance but also on whether it is physically recruited adjacent to the BCR. We speculate that B cells with preserved CD72 surface expression but spatially mislocated CD72 may be functionally equivalent to CD72- cell. Resolving this question is beyond the reach of flow cytometry, motivating our use of molecular proximity to characterise the nanoscale protein-protein proximity network on the RA B-cell surface[55]. Utilizing this technology, we found that in the CD19 complex, CD11c+ B cells had a significantly higher CD72-CD81 co-localization than in another B cell subset. Interestingly, this phenomenon was also more pronounced in RA B cells than in healthy control B cells.

Consistent with a reported higher antigen-presenting capacity in CD11c+ CD21- B cells [19], our study found that this population exhibits an altered CD40 centric network. CD40 is core components of the B-cell side of the immunological synapse [56]. The spatial proximity level of these molecules around MHC class II further supports a model in which the CD11c^high^ CD21^low^ population being primed for interaction with T cells[19]. In addition, we observed increased CD44-CD59, CD44-CD52 and CD44-CD55 co-localization in the CD11c^high^ CD21^low^ B cell subset. As CD59 and CD55 are both complement regulatory proteins that inhibit membrane attack complex formation, their proximity to CD44 in this subset raises the question of whether complement regulation is spatially organized at the B cell surface.

Of clinical relevance, we found substantial differences in the synovial B cell phenotype in patients that had received methotrexate TNF blockade combination therapy compared to those that were not receiving DMARDs at the time of sampling. Hence, treatment may reshape the SF B cell compartment. Differences included a reduction in the CD11c+ CD21- fraction while increasing CD11c+ CD21+. Although we did not have access to baseline SF samples before treatment, our results are consistent with recent longitudinal data from peripheral blood of newly diagnosed RA showing MTX-mediated immune remodelling with reduced plasmablasts and Tfh cells[57]. Yet, our data contrasts with a previous study showing stable RA blood B cell phenotypes after TNFi [58]. All the investigated patients in our study still had active synovitis that required arthrocentesis. Importantly, treatment status and serostatus were not independent in our cohort, as most ACPA+ patients were untreated while most ACPA- patients received MTX/TNFi; the contributions of serostatus and treatment to the synovial B cell phenotype therefore cannot be separated in this cross-sectional design.

Several limitations should be acknowledged. PNA-based analysis was performed on peripheral blood rather than synovial fluid, as insufficient B cell numbers could be recovered from synovial fluid sample. Hence the composition of the CD11c+ CD21- could be different from what was observed in the blood. Moreover, DN2 cells could not be resolved on the PNA UMAP although identified through manual “gating”. In addition, the pilot reagent panel lacked a correctly performing CD79a antibody and the current panel does not include anti-IgG/IgA reagents, restricting BCR complex analysis. The samples size in this study was modest, and validation in larger, independent RA cohorts will be needed to confirm these findings.

Taken together, this study identifies CD11c+ CD21- B cells as a shared but disease differentiated phenotype between ACPA+RA, ACPA-RA and SpA. ACPA+RA marked by a CD11c+ CD27- extrafollicular phenotype, treatment responsive remodelling, and divergent immature plasmablast states, antigen presenting phenotypes and T cell signatures. Our receptor proximity data further link this phenotype to altered CD72-BCR colocalization, a CD44 centric network and MHCII colocalization, providing a molecular basis for the hyperresponsive state of CD11c+ B cell in RA.

## Acknowledgement

We thank all patients with RA that contributed to these research efforts. We also wish to thank Lucymary Okechukwu and Julia Norkko, for managing the cohort biobanking and handling of blood samples. We thank Annika van Vollenhoven and the Center for Molecular Medicine flow cytometry core facility for valuable technical advice regarding flow cytometry acquisition and analysis. We also are grateful to late Anca Catrina for cohort initiatives and sample collection. We thank Lars Klareskog for fruitful scientific discussion and the supportive research environment.

## Ethics statement

The study was conducted in accordance with the Declaration of Helsinki, with ethical approval granted from the Regional Ethics Review Board Stockholm, Sweden, and written informed consent from all study participants.

## Contributors

WH, VM and CG conceived and designed the study. WH, SG, KW and MN performed experiments. WH, SG, LF and SP contributed to formal analysis and investigation. VM and CG provided resources and supervision, with KC contributing additional clinal data of patients. WH drafted the original manuscript with CG; all authors (WH, SG, KW, MN, LF, SP, AH, KC, VM, CG) contributed on reviewing and editing the manuscript. WH, SG, LF, and SP performed data visualization. KC, VM, and CG acquired funding for this study.

## Data availability

Data are available from the corresponding author upon request.

## REFERENCES

1. Smolen, J.S., et al., Rheumatoid arthritis. Nat Rev Dis Primers, 2018. 4: p. 18001.

2. Raychaudhuri, S., et al., Five amino acids in three HLA proteins explain most of the association between MHC and seropositive rheumatoid arthritis. Nat Genet, 2012. 44(3): p. 291–6.

3. Edwards, J.C., et al., Efficacy of B-cell-targeted therapy with rituximab in patients with rheumatoid arthritis. N Engl J Med, 2004. 350(25): p. 2572–81.

4. Genovese, M.C., et al., Abatacept for rheumatoid arthritis refractory to tumor necrosis factor alpha inhibition. N Engl J Med, 2005. 353(11): p. 1114–23.

5. Bashford-Rogers, R.J.M., et al., Analysis of the B cell receptor repertoire in six immune-mediated diseases. Nature, 2019. 574(7776): p. 122–126.

6. Wang, Y., et al., Rheumatoid arthritis patients display B-cell dysregulation already in the naive repertoire consistent with defects in B-cell tolerance. Sci Rep, 2019. 9(1): p. 19995.

7. Cowan, G.J.M., et al., In Human Autoimmunity, a Substantial Component of the B Cell Repertoire Consists of Polyclonal, Barely Mutated IgG(+ve) B Cells. Front Immunol, 2020. 11: p. 395.

8. Huang, W., et al., Variable fab domain N-glycosylation patterns in the B cell receptor repertoires of healthy individuals and patients with rheumatoid arthritis. J Immunol, 2026. 215(6).

9. Kinslow, J.D., et al., Elevated IgA Plasmablast Levels in Subjects at Risk of Developing Rheumatoid Arthritis. Arthritis Rheumatol, 2016. 68(10): p. 2372–83.

10. Iwata, S., et al., Activation of Syk in peripheral blood B cells in patients with rheumatoid arthritis: a potential target for abatacept therapy. Arthritis Rheumatol, 2015. 67(1): p. 63–73.

11. Liubchenko, G.A., et al., Rheumatoid arthritis is associated with signaling alterations in naturally occurring autoreactive B-lymphocytes. J Autoimmun, 2013. 40: p. 111–21.

12. Neys, S.F.H., et al., Aberrant B cell receptor signaling in circulating naive and IgA(+) memory B cells from newly-diagnosed autoantibody-positive rheumatoid arthritis patients. J Autoimmun, 2024. 143: p. 103168.

13. Wangriatisak, K., et al., CD72 downregulation on DN2 B cells is associated with disease activity and resistance to rituximab in systemic lupus erythematosus. Rheumatology (Oxford), 2026. 65(3).

14. Thorarinsdottir, K., et al., CD21(-/low) B cells associate with joint damage in rheumatoid arthritis patients. Scand J Immunol, 2019. 90(2): p. e12792.

15. Bao, W., M. Xie, and Y. Ye, Age-associated B cells indicate disease activity in rheumatoid arthritis. Cell Immunol, 2022. 377: p. 104533.

16. Vidal-Pedrola, G., et al., Characterization of age-associated B cells in early drug-naive rheumatoid arthritis patients. Immunology, 2023. 168(4): p. 640–653.

17. Wing, E., et al., Double-negative-2 B cells are the major synovial plasma cell precursor in rheumatoid arthritis. Front Immunol, 2023. 14: p. 1241474.

18. Jenks, S.A., et al., Distinct Effector B Cells Induced by Unregulated Toll-like Receptor 7 Contribute to Pathogenic Responses in Systemic Lupus Erythematosus. Immunity, 2018. 49(4): p. 725–739 e6.

19. McGrath, S., et al., Correlation of Professional Antigen-Presenting Tbet(+)CD11c(+) B Cells With Bone Destruction in Untreated Rheumatoid Arthritis. Arthritis Rheumatol, 2024. 76(8): p. 1263–1277.

20. Aletaha, D., et al., 2010 rheumatoid arthritis classification criteria: an American College of Rheumatology/European League Against Rheumatism collaborative initiative. Ann Rheum Dis, 2010. 69(9): p. 1580-8.

21. Filip Karlsson, M.S., Christina Galonska, Max Karlsson, Hanna van Ooijen, Tomasz Kallas, Divya Thiagarajan, Maud Schweitzer, Ludvig Larsson, Vincent van Hoef, Pouria Tajvar, Johan Dahlberg, Florian De Temmerman, Louise Leijonancker, Sylvain Geny, Rikard Forlin, Erika Negrini, Stefan Petkov, Lovisa Franzén, Jessica Bunz, Christine Moge, Henrik Everberg, Petter Brodin, Alvaro Martinez Barrio, Simon Fredriksson, Single-cell protein interactomes by the Proximity Network Assay. bioRxiv, 2025.

22. Soh, K.T., et al., CD319 (SLAMF7) an alternative marker for detecting plasma cells in the presence of daratumumab or elotuzumab. Cytometry B Clin Cytom, 2021. 100(4): p. 497–508.

23. Pitzalis, C., et al., Ectopic lymphoid-like structures in infection, cancer and autoimmunity. Nat Rev Immunol, 2014. 14(7): p. 447–62.

24. Hardt, U., et al., Integrated single cell and spatial transcriptomics reveal autoreactive differentiated B cells in joints of early rheumatoid arthritis. Sci Rep, 2022. 12(1): p. 11876.

25. de Vries, C., et al., Rheumatoid Arthritis Related B-Cell Changes Are Found Already in the Risk-RA Phase. Eur J Immunol, 2025. 55(2): p. e202451391.

26. Shwetar JJ, e.a., Costimulatory blockade depletes T peripheral helper, late-activated naïve, and DN2 B cells in rheumatoid arthritis. medRxiv, 2026.

27. Eisenbarth, S.C., et al., A roadmap for defining “extrafollicular” B cell responses. Immunity, 2025. 58(11): p. 2627–2645.

28. Cancro, M.P., Age-Associated B Cells. Annu Rev Immunol, 2020. 38: p. 315–340.

29. Xiaohao Wu, M.Z., Jae-Seung Moon, Eun Kyung Song, Joshua C. Abrams, Laura S. van Dam, Orr Sharpe, Marie Feser, Laurie Moss, Peggy P. Ho, Melanie H. Smith, Laura T. Donlin, Jane H. Buckner, Eddie A. James, Gary S. Firestein, Yuko Okamoto, Tobiaz V. Lanz, Eric Meffre, V. Michael Holers, Kevin D. Deane, William H. Robinson, Senescent Activated Naive B Cells Promote Anti-Citrullinated Antigen T Cell Responses and the Transition to Clinical Rheumatoid Arthritis. BioRxiv, 2026.

30. Wang, T., et al., Evaluation of B cell related markers and autoantibodies in rheumatoid arthritis patients treated with abatacept. Front Immunol, 2025. 16: p. 1504454.

31. Shwetar, J.J., et al., Costimulatory blockade depletes T peripheral helper, late-activated naive, and DN2 B cells in rheumatoid arthritis. medRxiv, 2026.

32. Argyriou, A., et al., Single cell sequencing identifies clonally expanded synovial CD4(+) T(PH) cells expressing GPR56 in rheumatoid arthritis. Nat Commun, 2022. 13(1): p. 4046.

33. Zhang, F., et al., Defining inflammatory cell states in rheumatoid arthritis joint synovial tissues by integrating single-cell transcriptomics and mass cytometry. Nat Immunol, 2019. 20(7): p. 928–942.

34. Genovese, M.C., et al., Efficacy and safety of secukinumab in patients with rheumatoid arthritis: a phase II, dose-finding, double-blind, randomised, placebo controlled study. Ann Rheum Dis, 2013. 72(6): p. 863–9.

35. Postigo, A.A., et al., Regulated expression and function of CD11c/CD18 integrin on human B lymphocytes. Relation between attachment to fibrinogen and triggering of proliferation through CD11c/CD18. J Exp Med, 1991. 174(6): p. 1313–22.

36. Nagy-Balo, Z., et al., Activated Human Memory B Lymphocytes Use CR4 (CD11c/CD18) for Adhesion, Migration, and Proliferation. Front Immunol, 2020. 11: p. 565458.

37. Doorenspleet, M.E., et al., Rheumatoid arthritis synovial tissue harbours dominant B-cell and plasma-cell clones associated with autoreactivity. Ann Rheum Dis, 2014. 73(4): p. 756–62.

38. Elliott, S.E., et al., B cells in rheumatoid arthritis synovial tissues encode focused antibody repertoires that include antibodies that stimulate macrophage TNF-alpha production. Clin Immunol, 2020. 212: p. 108360.

39. Dunlap, G., et al., Clonal associations between lymphocyte subsets and functional states in rheumatoid arthritis synovium. Nat Commun, 2024. 15(1): p. 4991.

40. Humby, F., et al., Synovial cellular and molecular signatures stratify clinical response to csDMARD therapy and predict radiographic progression in early rheumatoid arthritis patients. Ann Rheum Dis, 2019. 78(6): p. 761–772.

41. Zhang, F., et al., Deconstruction of rheumatoid arthritis synovium defines inflammatory subtypes. Nature, 2023. 623(7987): p. 616–624.

42. Ueno, A., et al., The production of CXCR3-agonistic chemokines by synovial fibroblasts from patients with rheumatoid arthritis. Rheumatol Int, 2005. 25(5): p. 361–7.

43. Meiser, A., et al., The chemokine receptor CXCR3 is degraded following internalization and is replenished at the cell surface by de novo synthesis of receptor. J Immunol, 2008. 180(10): p. 6713–24.

44. Henneken, M., et al., Differential expression of chemokine receptors on peripheral blood B cells from patients with rheumatoid arthritis and systemic lupus erythematosus. Arthritis Res Ther, 2005. 7(5): p. R1001–13.

45. Pan, C., N. Baumgarth, and J.R. Parnes, CD72-deficient mice reveal nonredundant roles of CD72 in B cell development and activation. Immunity, 1999. 11(4): p. 495–506.

46. Tsubata, T., CD72 is a Negative Regulator of B Cell Responses to Nuclear Lupus Self-antigens and Development of Systemic Lupus Erythematosus. Immune Netw, 2019. 19(1): p. e1.

47. Adachi, T., et al., CD72 negatively regulates signaling through the antigen receptor of B cells. J Immunol, 2000. 164(3): p. 1223–9.

48. Wu, Y., et al., The B-cell transmembrane protein CD72 binds to and is an in vivo substrate of the protein tyrosine phosphatase SHP-1. Curr Biol, 1998. 8(18): p. 1009–17.

49. Kumanogoh, A., et al., Identification of CD72 as a lymphocyte receptor for the class IV semaphorin CD100: a novel mechanism for regulating B cell signaling. Immunity, 2000. 13(5): p. 621–31.

50. Li, D.H., et al., CD72 down-modulates BCR-induced signal transduction and diminishes survival in primary mature B lymphocytes. J Immunol, 2006. 176(9): p. 5321–8.

51. Kumanogoh, A., et al., Requirement for CD100-CD72 interactions in fine-tuning of B-cell antigen receptor signaling and homeostatic maintenance of the B-cell compartment. Int Immunol, 2005. 17(10): p. 1277–82.

52. Mattila, P.K., et al., The actin and tetraspanin networks organize receptor nanoclusters to regulate B cell receptor-mediated signaling. Immunity, 2013. 38(3): p. 461–74.

53. Depoil, D., et al., CD19 is essential for B cell activation by promoting B cell receptor-antigen microcluster formation in response to membrane-bound ligand. Nat Immunol, 2008. 9(1): p. 63–72.

54. Garcia-Parajo, M.F., et al., Nanoclustering as a dominant feature of plasma membrane organization. J Cell Sci, 2014. 127(Pt 23): p. 4995–5005.

55. Karlsson, F., et al., Molecular pixelation: spatial proteomics of single cells by sequencing. Nat Methods, 2024. 21(6): p. 1044–1052.

56. Cannons, J.L., et al., Optimal germinal center responses require a multistage T cell:B cell adhesion process involving integrins, SLAM-associated protein, and CD84. Immunity, 2010. 32(2): p. 253–65.

57. Preglej, T., et al., Time-resolved immune dynamics in rheumatoid arthritis under methotrexate therapy. Ann Rheum Dis, 2026.

58. Meednu, N., et al., Activated Peripheral Blood B Cells in Rheumatoid Arthritis and Their Relationship to Anti-Tumor Necrosis Factor Treatment and Response: A Randomized Clinical Trial of the Effects of Anti-Tumor Necrosis Factor on B Cells. Arthritis Rheumatol, 2022. 74(2): p. 200–211.

