## Supplemental Material for "Distinct synovial fluid B-cell differentiation and activation profiles in rheumatoid arthritis"

### SUPPORTING INFORMATION - METHODS

#### Generating flow cytometry based UMAP in R

Gated live CD19+CD3-CD4- or CD19-CD3+ events were exported and analysed in RStudio 4.6.1. FCS files acquired over seven days using the 31-marker B-cell panel or the 28-marker T-cell panel (Supplementary Table 2) were imported with the R package FlowCore. A daily technical control from one reference individual was used for batch diagnostics but excluded from biological comparisons. Intensities were arcsinh-transformed (cofactor 6000), and each file was downsampled to ≤5,000 cells before pooling.

Pooled intensities were z-scored and reduced to 20 principal components (prcomp). Inter-day batch effects were corrected using Harmony (v2.0.5) [1] with the acquisition day as a covariate (vars_use = "day"). Correction was confirmed on the technical control before/after Harmony. Harmony-corrected components were thereafter clustered using FlowSOM [2] (10 × 10 SOM, consensus meta-clustering): for B cells with the settings nClus = 15 for PB+SF and nClus = 10 for SF-only; for T cells with the settings nClus = 13 for PB+SF. UMAP visualization was generated before and after Harmony to demonstrate batch correction.

#### PNA data processing

Raw Proximity Network Assay (PNA) sequencing data were processed using the Pixelator workflow through the Nextflow-based nf-core/pixelator pipeline. Downstream analyses were performed in R using pixelatorR (v0.13.0) together with Seurat (v5.2.1), ggplot2 (v3.5.1), and ComplexHeatmap (v2.22.0).

Individual cellular components were subjected to quality-control filtering before downstream analysis. Components were retained if they had a normal Tau dispersion classification, at least 20,000 unique molecular identifiers (UMIs), an isotype-control fraction below 0.1% (isotype_fraction < 0.001), and a dangling-node fraction (k-core 1 fraction) below 0.5. These criteria were used to remove components with low molecular coverage, excessive nonspecific antibody signal, abnormal marker dispersion, or insufficient network connectivity.

For cell-surface protein abundance analysis, protein counts were normalized using centered log-ratio (CLR) normalization per cell. Dimensionality reduction and unsupervised clustering were performed using the first 18 Harmony-corrected principal components. A shared nearest-neighbor graph was constructed using these components, followed by Louvain clustering at a resolution of 0.8, yielding eight B-cell clusters. B-cell populations were subsequently characterized using canonical surface-marker expression and compared with B-cell populations identified by conventional IgD/CD27-based gating.

Protein-proximity analyses were performed separately from protein-abundance analyses. Background proximity signal was removed using FilterProximityScores with a background_threshold_pct of 0.00071 and a minimum detection requirement of five cells per protein pair. For subsequent proximity analyses, protein-pair observations were retained only when the counts of both constituent markers exceeded 100 and the corresponding join count exceeded 5. Proximity was quantified using the log2-ratio metric. Summarized proximity scores were calculated using SummarizeProximityScores with the mean log2 ratio as the summary statistic and with missing observations retained during summarization. Scores were summarized at the B-cell population, donor, or condition level according to the analysis.

For visualization of protein-proximity architecture, summarized mean log2-ratio values were used to generate pairwise proximity heatmaps and network representations. Positive log2-ratio values indicate enrichment of the observed protein-pair proximity relative to its expected abundance-based background, whereas negative values indicate relative spatial segregation. The fraction of cells in which each protein pair was detected was retained as an additional measure of detection prevalence.

**Supplementary Table 1. Patient characteristics**

| Group | Patients | Age | Sex | Anti-CCP2 | RF | DMARDs at time of analysis | Samples available for flow cytometry | Flow Panels applied for PB | Flow Panel applied for SF | Pixelgen |
|---|---|---|---|---|---|---|---|---|---|---|
| ACPA+ RA | 1 | 50-55 | M | pos | pos | N | SF/PB | FC B/T | FC B/T | |
| | 2 | 35-40 | M | pos | pos | MTX + Etanercept | SF | N | FC B/T/phosflow | |
| | 3 | 55-60 | F | pos | pos | MTX + Adalimumab | SF/PB | FC B/T/phosflow | FC B/T/phosflow | |
| | 4 | 50-55 | M | pos | pos | MTX | SF/PB | FC B/T/phosflow | FC B/T/phosflow | |
| | 5 | 35-40 | F | pos | pos | MTX + Infliximab | SF/PB | FC B/T/phosflow | FC B/T/phosflow | |
| | 6 | 60-65 | F | pos | pos | N | SF | N | FC B/T/phosflow | |
| | 7 | 50-55 | F | pos | pos | N | SF/PB | FC B/T/phosflow | FC B/T/phosflow | |
| | 8 | 45-50 | F | pos | pos | N | SF/PB | FC B | FC B | |
| | 9 | 75-80 | M | pos | pos | N | SF | N | FC B/T | |
| | 10 | 70-75 | F | pos | pos | N | SF/PB | FC B/T/phosflow | FC B/T | |
| | 11 | 45-50 | M | pos | neg | Etanercept + Salazopyrine | SF/PB | FC B/T/phosflow | FC B/T/phosflow | |
| | 12 | 60-65 | F | pos | neg | N | SF/PB | FC B/T/phosflow | FC B/T/phosflow | |
| ACPA- RA | 1 | 25-30 | F | neg | neg | MTX + Salazopyrine | SF/PB | FC B/T | FC B/T | |
| | 2 | 65-70 | F | neg | neg | MTX | SF/PB | FC B/T/phosflow | FC B/T/phosflow | |
| | 3 | 40-45 | F | neg | neg | Etanercept | SF/PB | FC B/T/phosflow | FC B/T/phosflow | |
| | 4 | 55-60 | F | neg | neg | MTX | SF/PB | FC B/T | FC B/T/phosflow | |
| | 5 | 30-35 | F | neg | neg | MTX | SF/PB | FC B/T/phosflow | FC B/T | |
| | 6 | 30-35 | F | neg | neg | MTX + Golimumab | SF/PB | FC B/T | FC B/T | |
| | 7 | 60-65 | F | neg | neg | MTX + Adalimumab | SF/PB | FC B/T/phosflow | FC B/T/phosflow | |
| | 8 | 55-60 | F | neg | neg | Salazopyrine | SF/PB | FC B/T/phosflow | FC B/T/phosflow | |
| | 9 | 35-40 | F | neg | neg | Certolizumab pegol | SF/PB | FC B/T/phosflow | FC B/T/phosflow | |
| | 10 | 55-60 | F | neg | neg | N | SF | FC B/T/phosflow | FC B/T | |
| SpA | 1 | 45-50 | F | neg | neg | N | SF | N | FC B/T/phosflow | |
| | 2 | 55-60 | F | neg | neg | N | SF/PB | FC B/T | FC B/T | |
| | 3 | 30-35 | F | neg | neg | Etanercept | SF/PB | FC B/T | FC B/T | |
| | 4 | 55-60 | F | neg | neg | N | SF/PB | FC B/T/phosflow | FC B/T/phosflow | |
| | 5 | 45-50 | F | neg | neg | MTX + Etanercept | SF/PB | FC B/T/phosflow | FC B/T/phosflow | |
| | 6 | 50-55 | F | md | md | Azathioprine | SF/PB | FC B/T/phosflow | FC B/T | |
| | 7 | 25-30 | M | neg | neg | MTX | SF/PB | FC B/T/phosflow | FC B/T/phosflow | |
| | 8 | 55-60 | M | neg | neg | N | SF/PB | FC B/T/phosflow | FC B/T/phosflow | |
| ACPA+ RA | 13# | 45-50 | F | pos | pos | N | PB | FC B | | PBMC & B cells |
| | 14# | 35-40 | F | pos | neg | N | PB | FC B | | PBMC & B cells |
| | 15# | 50-55 | F | pos | neg | N | PB | FC B | | PBMC & B cells |
| HC | 1# | 45-50 | F | neg | neg | N | PB | FC B | | B cells |
| | 2# | 35-40 | F | neg | neg | N | PB | FC B | | B cells |

#: Enrichment of B cell by negative selection; CCP2: cyclic citrullinated peptide; RF: rheumatoid factor N: no; Y: yes; neg: negative; pos: positive; MTX: methotrexate; FC: flow cytometry; PB: peripheral blood; SF: synovial fluid; md= missing data

**Supplementary Table 2. Antibodies panel used in spectral flow cytometry analysis**

***Antibodies used for B cell phenotype***

| Antibody | Fluorophore | Dilution | Comapny | Catalogue No. | Clone |
|---|---|---|---|---|---|
| CD24 | BUV496 | 1:25 | BD (OptiBuild) | 741143 | ML5 |
| CD43 | NovaFluor Blue585 | 1:25 | Thermo Fisher Scientific | H069T03B04-A | eBio84-3C1 |
| CD14 | eFluor 506 | 1:25 | Thermo Fisher Scientific | 69-0149-42 | 61D3 |
| CD19 | BV570 | 1:25 | BioLegend | 302236 | HIB19 |
| CXCR5 | BV750 | 1:25 | BioLegend | 356942 | J252D4 |
| CD319 | BUV737 | 1:25 | BD (OptiBuild) | 750833 | 235614 |
| CCR7 | PE-Fire 810 | 1:50 | BioLegend | 353269 | G043H7 |
| CD86 | BB515 | 1:50 | BD | 564545 | FUN-1, 2331 |
| CD20 | BUV395 | 1:50 | BD | 563781 | 2H7 |
| IgG | BV421 | 1:50 | BD | 562581 | G18-145 |
| IgM | Super Bright 436 | 1:50 | Thermo Fisher Scientific | 62-9998-42 | SA-DA4 |
| CD10 | PE-CF594 | 1:50 | BD | 562396 | HI10a |
| CD37 | BV711 | 1:100 | BD (OptiBuild) | 742401 | M-B371 |
| BTLA | AlexaFluor647 | 1:100 | BioLegend | 344519 | MIH26 |
| CXCR3 | RB705 | 1:100 | BD | 570554 | 1C6 |
| CD71 | RB744 | 1:100 | BD (OptiBuild) | 757854 | M-A712 |
| CD73 | BV785 | 1:100 | BioLegend | 344028 | AD-2 |
| CD27 | APC-Fire 810 | 1:100 | BioLegend | 393214 | QA17A18 |
| CD3 | BUV661 | 1:100 | Thermo Fisher Scientific | 376-0036-41 | SK7 |
| TIGIT | RB780 | 1:200 | BD | 569940 | TgMab-2 |
| CD39 | PE-Fire 744 | 1:200 | BioLegend | 328253 | A1 |
| IgD | BV480 | 1:200 | BD | 566187 | IA6-2 |
| CD72 | RB545 | 1:200 | BD (OptiBuild) | 756280 | J4-117 |
| CD1c | BV605 | 1:200 | BioLegend | 331537 | L161 |
| CD4 | cFluor B532 | 1:200 | Cytek | R7-20038 | SK-3 |
| CD21 | BUV805 | 1:200 | BD (OptiBuild) | 742008 | B-Ly4 |
| IgA | VioBlue | 1:400 | Miltenyi Biotec | 130-113-479 | IS11-8E10 |
| CD38 | BUV563 | 1:400 | BD (OptiBuild) | 741446 | HB7 |
| HLA-DR | BV650 | 1:400 | BioLegend | 307650 | L243 |
| CD11c | APC-Fire 750 | 1:400 | BioLegend | 371509 | SHCL-3 |
| CD45RB | BUV615 | 1:800 | BD (OptiBuild) | 751482 | MT4 |
| CD95 | PECy5 | 1:800 | BD | 561977 | DX2 |
| Fixable viability dye | eFluor 506 | 1 to 1000 | Thermo Fisher Scientific | 65-0866-14 | |

***Antibodies used for T cell phenotype***

| | Marker | Fluorophore | Dilution | Company | Catalogue No. | Clone |
|---|---|---|---|---|---|---|
| Surface | CXCR3 | APC/FIRE810 | 1 to 25 | Biolegend | 353762 | G025H7 |
| | CCR6 | BUV661 | 1 to 25 | BD | 750696 | 11A9 |
| | CD14 | eFluor506 | 1 to 25 | Thermo Fisher Scientific | 69-0149-42 | 61D3 |
| | CD19 | eFluor506 | 1 to 25 | Thermo Fisher Scientific | 69-0199-42 | HIB19 |
| | CD3 | Spark blue 550 | 1 to 50 | Biolegend | 344852 | SK7 |
| | PD-1 | SB436 | 1 to 50 | ThermoFisher | 62-2799-42 | eBioJ105 |
| | CXCR5 | PE/Cy7 | 1 to 50 | Biolegend | 356923 | J252D4 |
| | ICOS | PE | 1 to 50 | BD | 557802 | DX29 |
| | CD127 | Pacific blue | 1 to 50 | Biolegend | 351306 | A019D5 |
| | CD27 | BUV496 | 1 to 50 | BD | 751678 | O323 |
| | CCR4 | BV605 | 1 to 50 | BD | 562906 | 1G1 |
| | CX3CR1 | BUV805 | 1 to 50 | BD | 749353 | EA |
| | CCR7 | PE/FIRE810 | 1 to 50 | Biolegend | 353269 | G043H7 |
| | CD4 | BV786 | 1 to 100 | BD | 563877 | SK3 |
| | GPR56 | BUV395 | 1 to 100 | BD | 752707 | CG4.rMAb |
| | CD25 | BV480 | 1 to 100 | BD | 567488 | BC96 |
| | CD45Ra | AF700 | 1 to 100 | Biolegend | 304120 | HI100 |
| | TCRgd | BV711 | 1 to 100 | BD | 568490 | 11F2 |
| | CD69 | BV750 | 1 to 100 | BD | 747522 | FN50 |
| | CD137 | PE/Cy5 | 1 to 100 | Biolegend | 309808 | 4B41 |
| | CD40L | BUV737 | 1 to 100 | BD | 748983 | TRAP1 |
| | TIGIT | RB780 | 1 to 200 | BD | 569940 | TgMab-2 |
| | CCR2 | RB705 | 1 to 200 | BD | 757633 | LS132.1D9 |
| | CD8 a +b | APC/FIRE750 | 1 to 400 | Biolegend | 344745 | SK1 |
| | CD38 | BUV563 | 1 to 400 | BD | 741446 | HB7 |
| | HLA-DR | BV650 | 1 to 400 | Biolegend | 307650 | L243 |
| | BV421 | LAG-3 | 1 to 25 | BD | 740072 | C9B7W |
| Intracellular | GZMK | AF647 | 1 to 50 | Biolegend | 370503 | GM26E7 |
| | GZMB | FITC | 1 to 50 | Biolegend | 372205 | QA18A28 |
| | FOXP3 | PE/Dazzle594 | 1 to 33 | Biolegend | 320125 | 206D |
| | Fixable viability dye | eFluor 506 | 1 to 1000 | Thermo Fisher Scientific | 65-0866-14 | |

***Antibodies panel of intracellular phosflow staining***

| | Marker | Fluorophore | Dilution | Company | Catalogue No. | Clone |
|---|---|---|---|---|---|---|
| Surface | CXCR5 | BV750 | 1 to 25 | BioLegend | 356942 | J252D4 |
| | CD19 | BV570 | 1 to 25 | BioLegend | 302236 | HIB19 |
| | CD24 | BUV496 | 1 to 25 | BD (OptiBuild) | 741143 | ML5 |
| | CD138 | BV711 | 1 to 25 | BD | 563184 | MI15 |
| | CD3 | efluor 506 | 1 to 25 | Invitrogen | 69-0038-42 | UCHT1 |
| | CD14 | efluor 506 | 1 to 25 | Thermo Fisher Scientific | 69-0149-42 | 61D3 |
| | CXCR4 | BUV395 | 1 to 50 | BD | 563924 | 12G5 |
| | BCMA | APC | 1 to 50 | Biolegend | 357505 | 19F2 |
| | CD27 | APC-Fire 810 | 1 to 100 | BioLegend | 393214 | QA17A18 |
| | IgD | BV480 | 1 to 200 | BD | 566187 | IA6-2 |
| | CD21 | BUV805 | 1 to 200 | BD (OptiBuild) | 742008 | B-Ly4 |
| | CD11c | APC-Fire 750 | 1 to 400 | BioLegend | 371509 | SHCL-3 |
| | CD38 | BUV563 | 1 to 400 | BD (OptiBuild) | 741446 | HB7 |
| Intracellular | pERK | AF647 | 1 to 25 | BD | 612593 | 20A |
| | pSYK | PE | 1 to 25 | BD | 558529 | I120-722 |
| | Fixable viability dye | eFluor 506 | 1 to 1000 | Thermo Fisher Scientific | 65-0866-14 | |

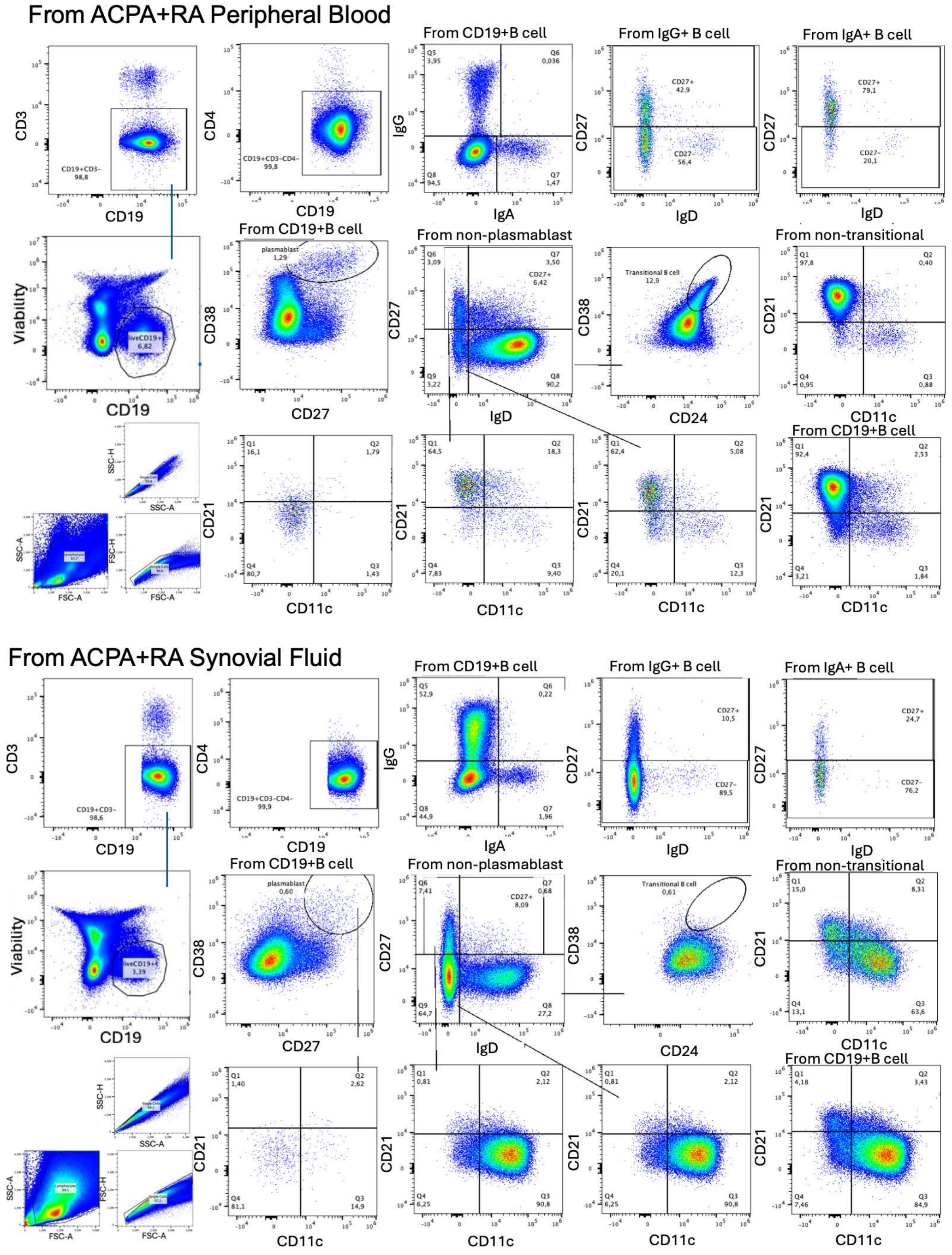


**Supplementary Fig. 1 | Gating strategy for B-cell subsets in PB and SF.**

Representative gating of B-cell subsets in peripheral blood (upper panels) and synovial fluid (lower panels) from a patient with ACPA+ RA. Plasmablasts were defined as CD19+ $CD27^{hi}CD38^{hi}$. Among the non-plasmablast B cells transitional B cells (CD27 -IgD- CD24+ CD38+), switched memory (SWM; CD27+IgD-), unswitched memory (USM; CD27+ IgD+) and double-negative (DN; CD27 -IgD-) were identified. Non-transitional B cells were subdivided into naïve B cells (NAV; CD27- IgD+). DN and NAV subsets were classified as DN1 (CD21+CD11c-), DN2 (CD21- CD11c+), DN3 (CD21- CD11c-), resting naïve (rNAV; CD21+CD11c-) and activated naïve (aNAV; CD21- CD11c+). IgA+ (CD19+ IgA+ IgG-) and IgG+ (CD19+ IgA-IgG+) B cells were analysed within the CD27+ and CD27- compartments.

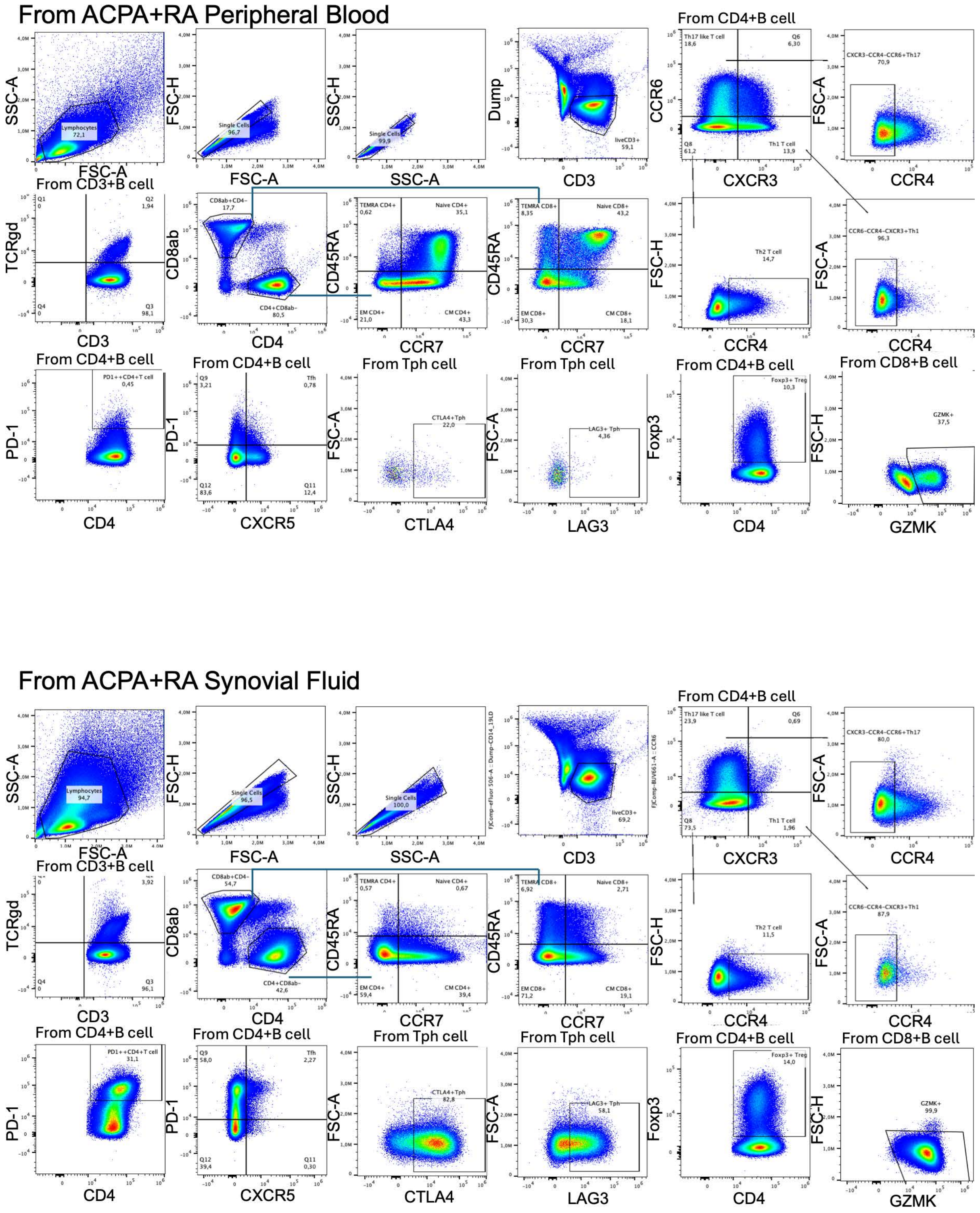


**Supplementary Fig. 2 | Gating strategy for T-cell subsets in PB and SF.**

Representative gating of T-cell subsets in peripheral blood (upper panels) and synovial fluid (lower panels) from a patient with ACPA+ RA. After lymphocyte and single-cell gating, live CD3+ T cells were gated into CD3+ TCRαβ- cells which further gated into CD4+ and CD8αβ+ T cells; memory and naïve T subsets were defined by CD45RA/CCR7 in CD4+ or CD8αβ+ T cell, T-helper lineages by CXCR3/CCR6/CCR4 (Th1, Th2, Th17), and follicular/peripheral helper cells by PD-1/CXCR5. CTLA4, LAG3 Tph (PD1++ CD4+ T cell), Foxp3 (Treg) and GZMK gates are shown. Comparable gates are shown for PB and SF.

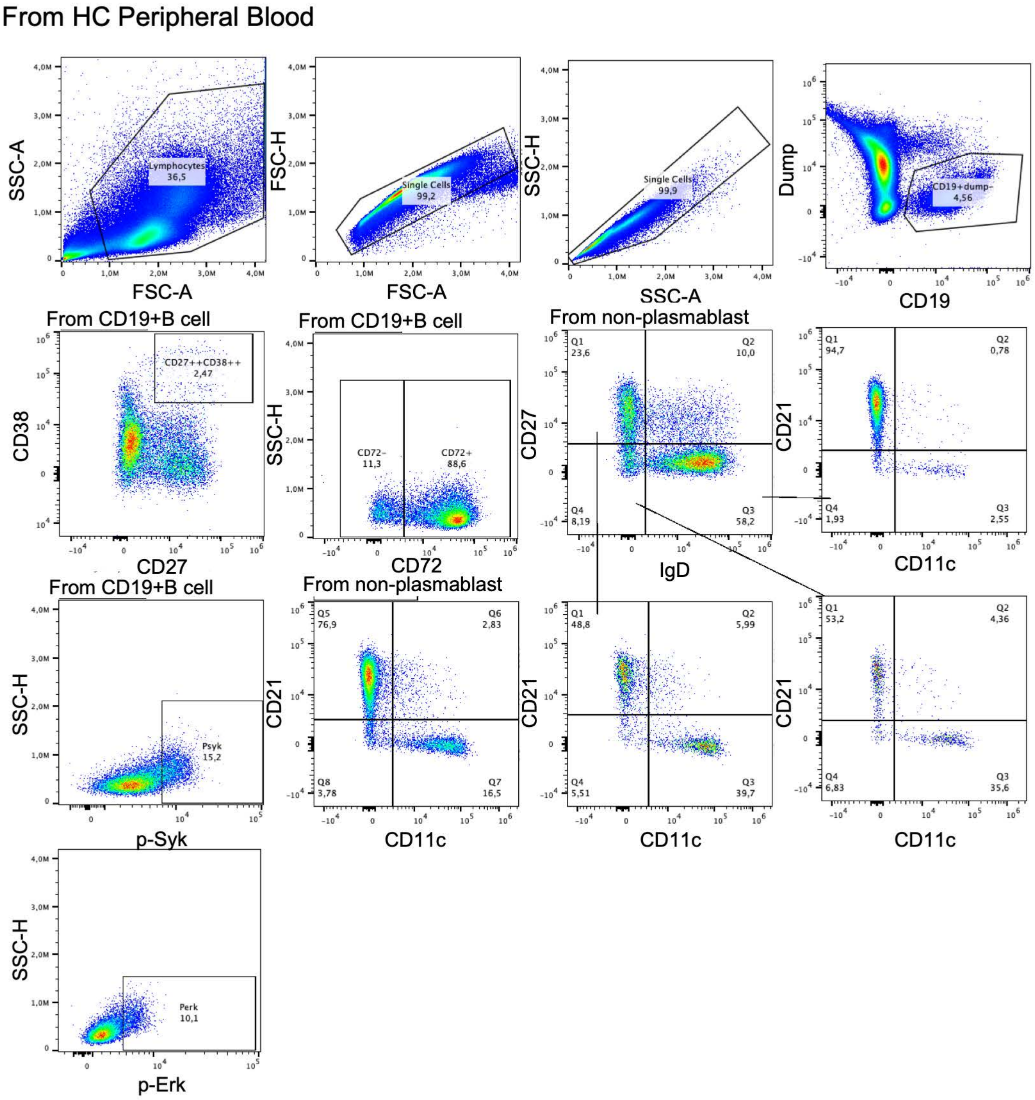


**Supplementary Fig. 3 | Gating strategy for phospho-flow analysis of B cells.**

Representative phospho-flow gating in PBMCs from a healthy control. After lymphocyte and single-cell gating, CD19+ (dump-) B cells were gated, plasmablasts excluded (CD27/CD38), and CD72+ versus CD72- were gated. Non-plasmablast B cells were further gated on CD21 versus CD11c to define the CD11c+CD21- population, and phospho-SYK (p-Syk) and phospho-ERK (p-Erk) gates were set on stimulated cells.

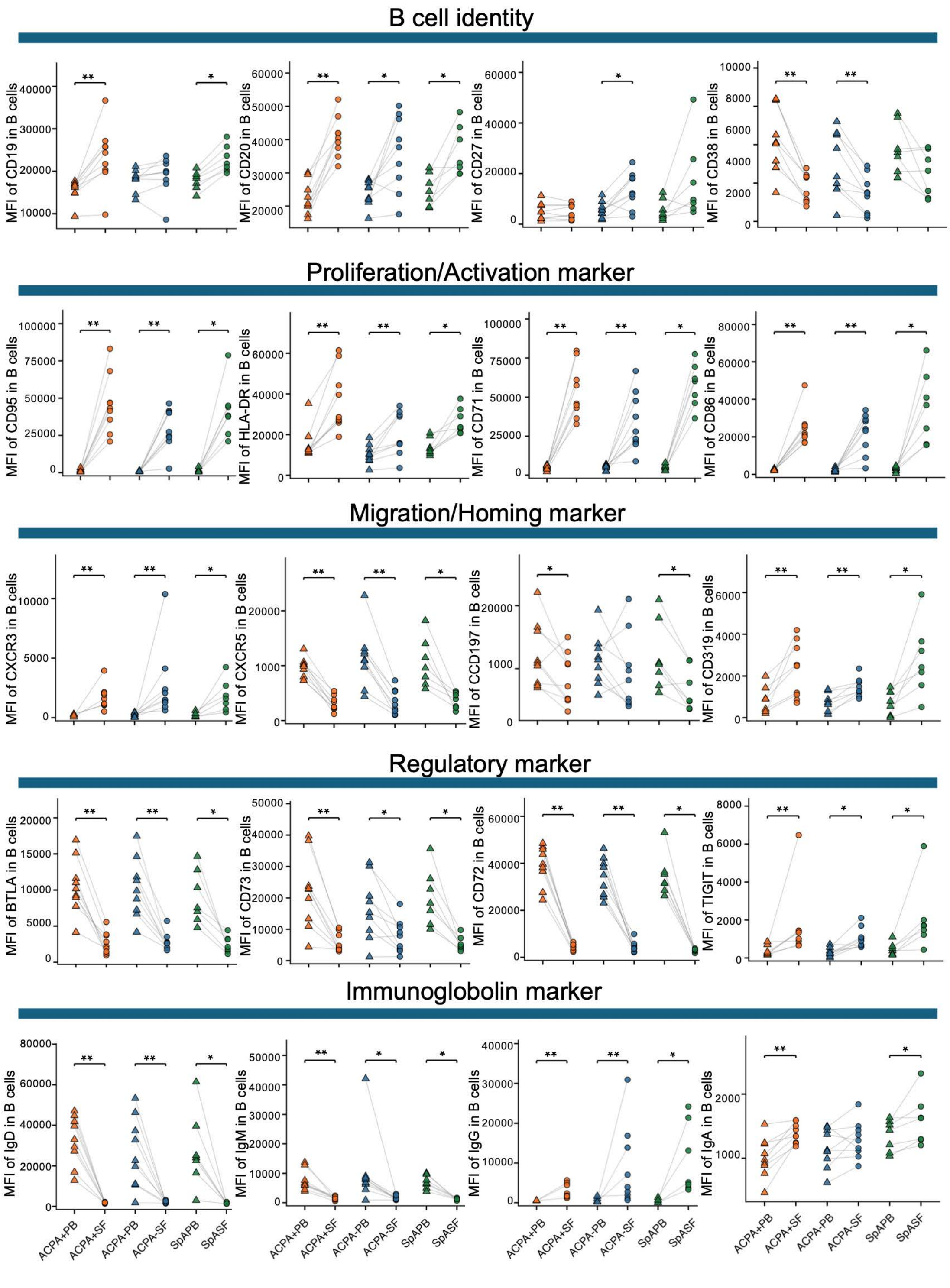


**Supplementary Fig. 4a | Surface-marker MFI on total B cells in paired blood and synovial fluid**

MFI of the indicated markers on total CD19+ B cells from paired PB and SF of patients with ACPA+ RA, ACPA- RA and SpA, grouped by functional category: B-cell identity (CD19, CD20, CD27, CD38), proliferation/activation (CD95, HLA-DR, CD71, CD86), migration/homing (CXCR3, CXCR5, CCR7, CD319), regulatory (BTLA, CD73, CD72, TIGIT) and immunoglobulin (IgD, IgM, IgG, IgA). Each symbol represents one patient; paired PB and SF samples are connected by grey lines. Two-tailed Wilcoxon matched-pairs signed-rank test was applied. *P < 0.05, **P < 0.01, ***P < 0.001.

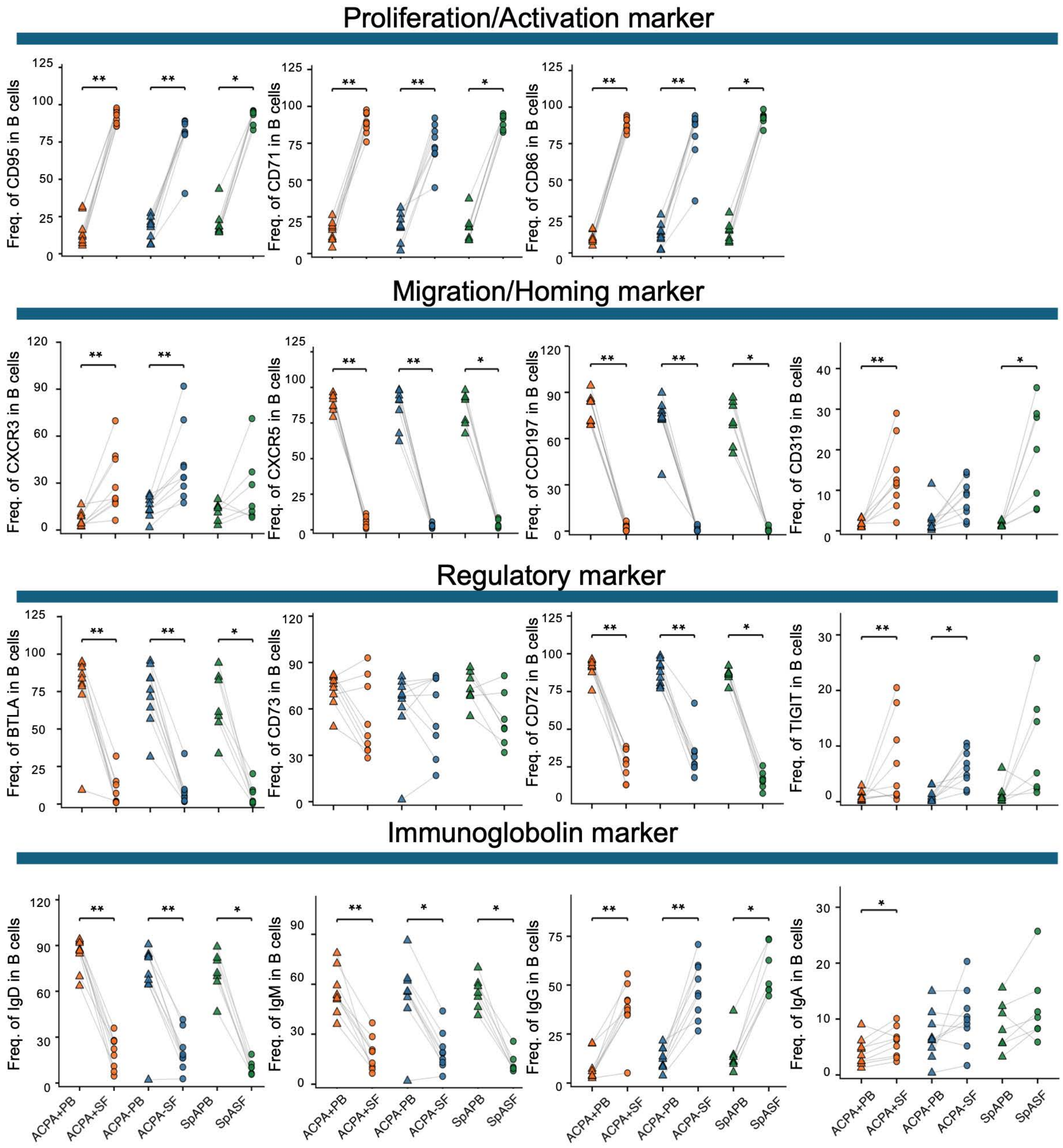


**Supplementary Fig. 4b | Surface-marker Frequency on total B cells in paired blood and synovial fluid**

Frequency of the indicated markers on total CD19+ B cells from paired PB and SF of patients with ACPA+ RA, ACPA- RA and SpA, grouped by functional category: proliferation/activation (CD95, CD71, CD86), migration/homing (CXCR3, CXCR5, CCR7, CD319), regulatory (BTLA, CD73, CD72, TIGIT) and immunoglobulin (IgD, IgM, IgG, IgA). Each symbol represents one patient; paired PB and SF samples are connected by grey lines. Two-tailed Wilcoxon matched-pairs signed-rank test was applied. *P < 0.05, **P < 0.01, ***P < 0.001.

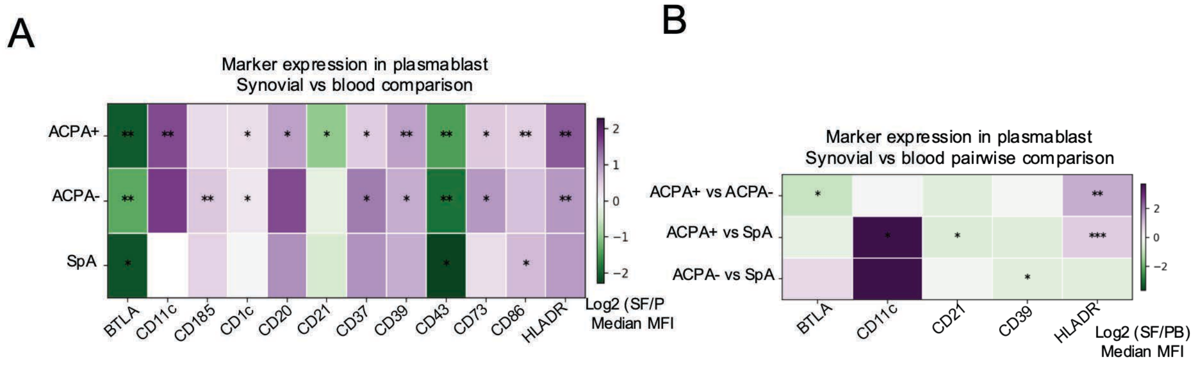


**Supplementary Fig. 5 | Differential marker expression in plasmablast among ACPA+ ACPA- and SpA**

**A**. Heatmap of the $\log_2$ fold change (SF/PB) of MFI for plasmablast surface markers in each disease group, restricted to markers with at least one significant SF-versus-PB comparison. **B**. Heatmap of the $\log_2$ MFI ratio between disease groups (ACPA+ vs ACPA-, ACPA+ vs SpA, ACPA- vs SpA) for plasmablast markers in SF, restricted to markers with at least one significant pairwise comparison. Two-tailed Mann-Whitney U test (A) and Wilcoxon signed-rank test (B). *P < 0.05, **P < 0.01, ***P < 0.001; ns, not significant.

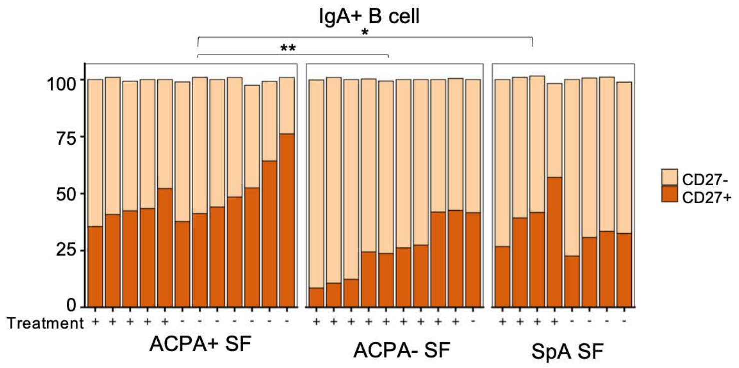


**Supplementary Fig. 6 | CD27+/CD27- proportions within IgA+ B cells among ACPA+ ACPA- and SpA SF**

Stacked bar graphs of CD27+/CD27- proportions within IgA+ B cells per SF sample, grouped by disease group and annotated by treatment status. Two-tailed Mann-Whitney U test. *P < 0.05, **P < 0.01, ***P < 0.001; ns, not significant.

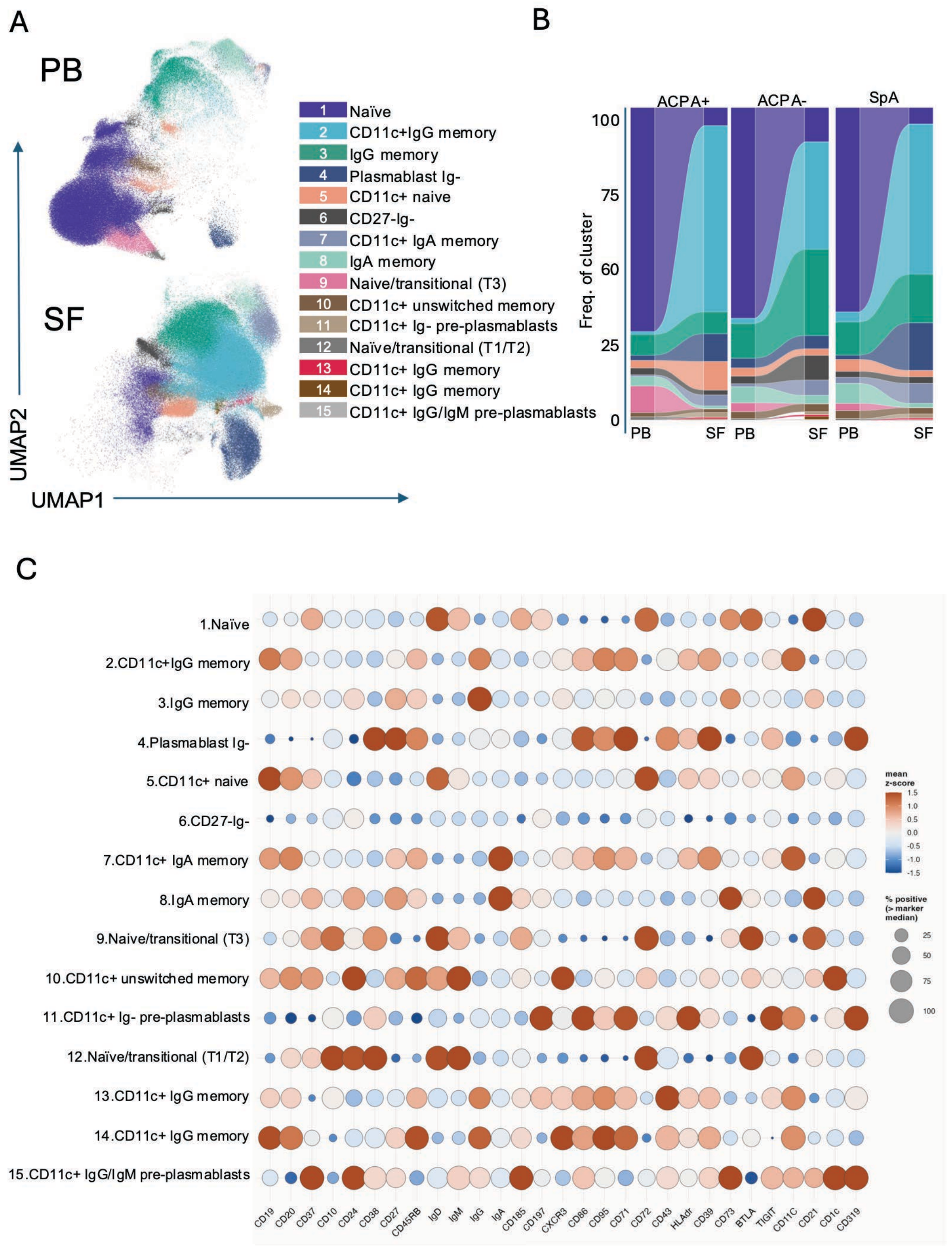


**Supplementary Fig. 7 | Unsupervised clustering of B cells from PB and SF**

**A**. UMAP embedding of CD19+ B cells split by origin (PB, SF), colored by fifteen unsupervised clusters (1, naïve; 2, CD11c+ IgG memory; 3, IgG memory; 4, plasmablast Ig-; 5, CD11c+ naïve; 6, CD27-Ig-; 7, CD11c+ IgA memory; 8, IgA memory; 9, naïve/transitional (T3); 10, CD11c+ unswitched memory; 11, CD11c+ Ig- pre-plasmablasts; 12, naïve/transitional (T1/T2); 13, CD11c+ IgG memory; 14, CD11c+ IgG memory; 15, CD11c+ IgG/IgM pre-plasmablasts). **B**. Ribbon plots showing the proportional distribution of clusters between paired PB and SF within each disease group (ACPA+, ACPA-, SpA). (C) Dot-plot heatmap of marker expression across the fifteen clusters; dot color encodes the mean z-scored expression and dot size the frequency of marker-positive cells.

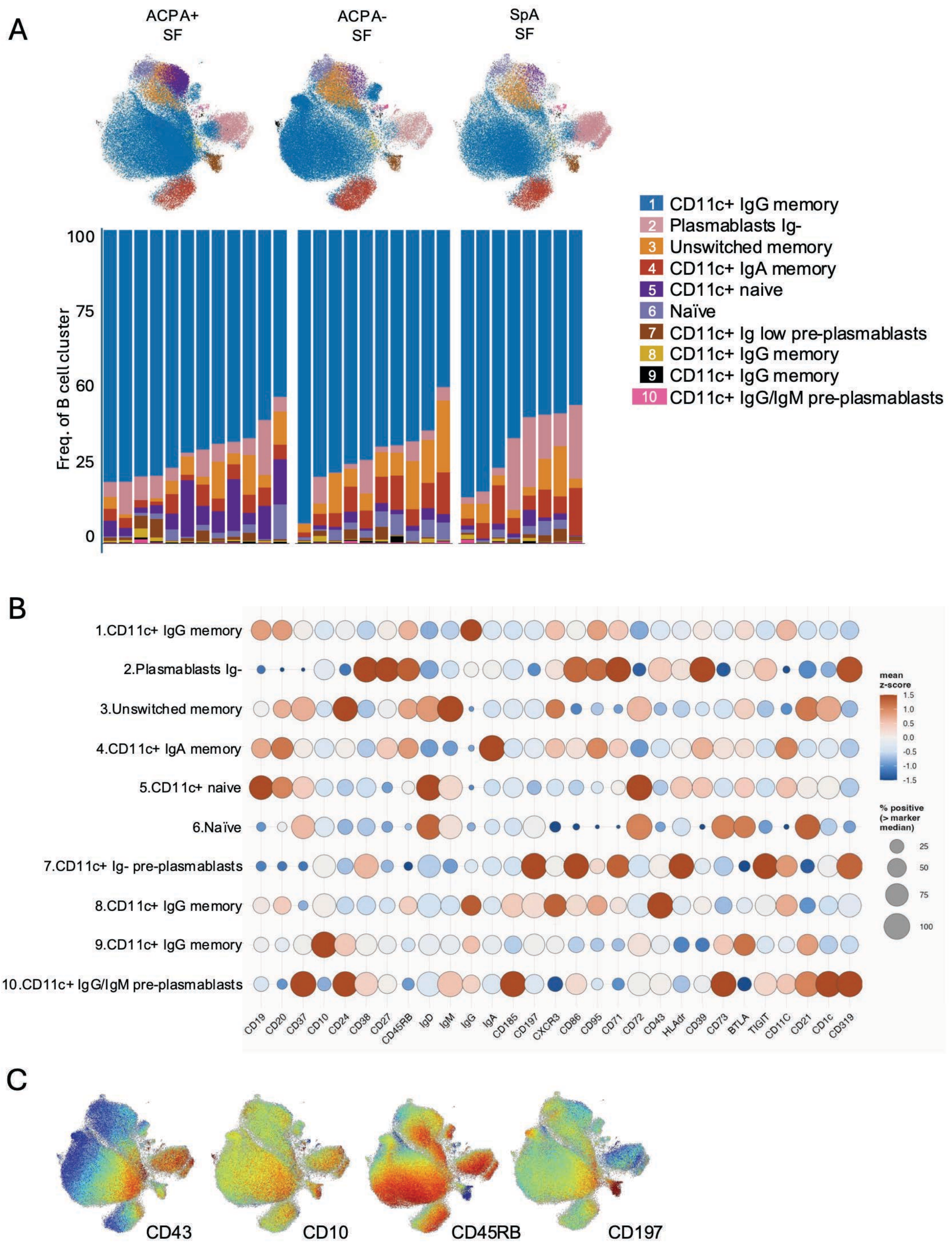


**Supplementary Fig. 8 | Unsupervised clustering of B cells from synovial fluid.**

**A.** UMAP embedding of CD19+ B cells from SF, colored by the ten unsupervised clusters. The same UMAP with cells from each disease group (ACPA+ RA, ACPA- RA and SpA) highlighted separately. Stacked bar graphs showing the proportional distribution of the ten clusters in each individual patient, grouped by disease group. **B.** Dot-plot heatmap of marker expression across the ten clusters; dot color encodes the mean z-scored expression and dot size the frequency of marker-positive cells. **C**. Selected Marker layer in UMAP.

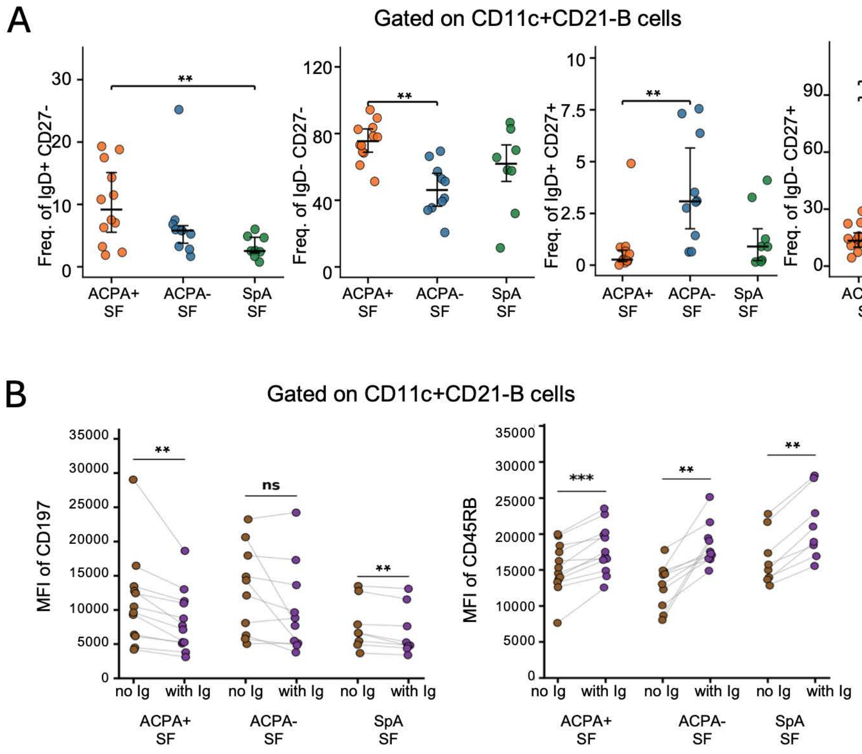


**Supplementary Fig. 9 | Differential marker expression between groups in CD11c+CD21-**

**A.** Graphs show the frequency of IgD/CD27-defined subpopulations (IgD+ CD27-, IgD- CD27-, IgD+ CD27+, IgD- CD27+) within CD11c+ CD21- B cells between SF disease groups. **B.** Graphs show MFI of CD197 on CD11c+ CD21- B cells lacking versus expressing surface Ig, in paired comparison within each SF group. Two-tailed Mann-Whitney U test (A) and Wilcoxon signed-rank test (B) were used for statistical analysis. *P < 0.05, **P < 0.01, ***P < 0.001; ns, not significant.

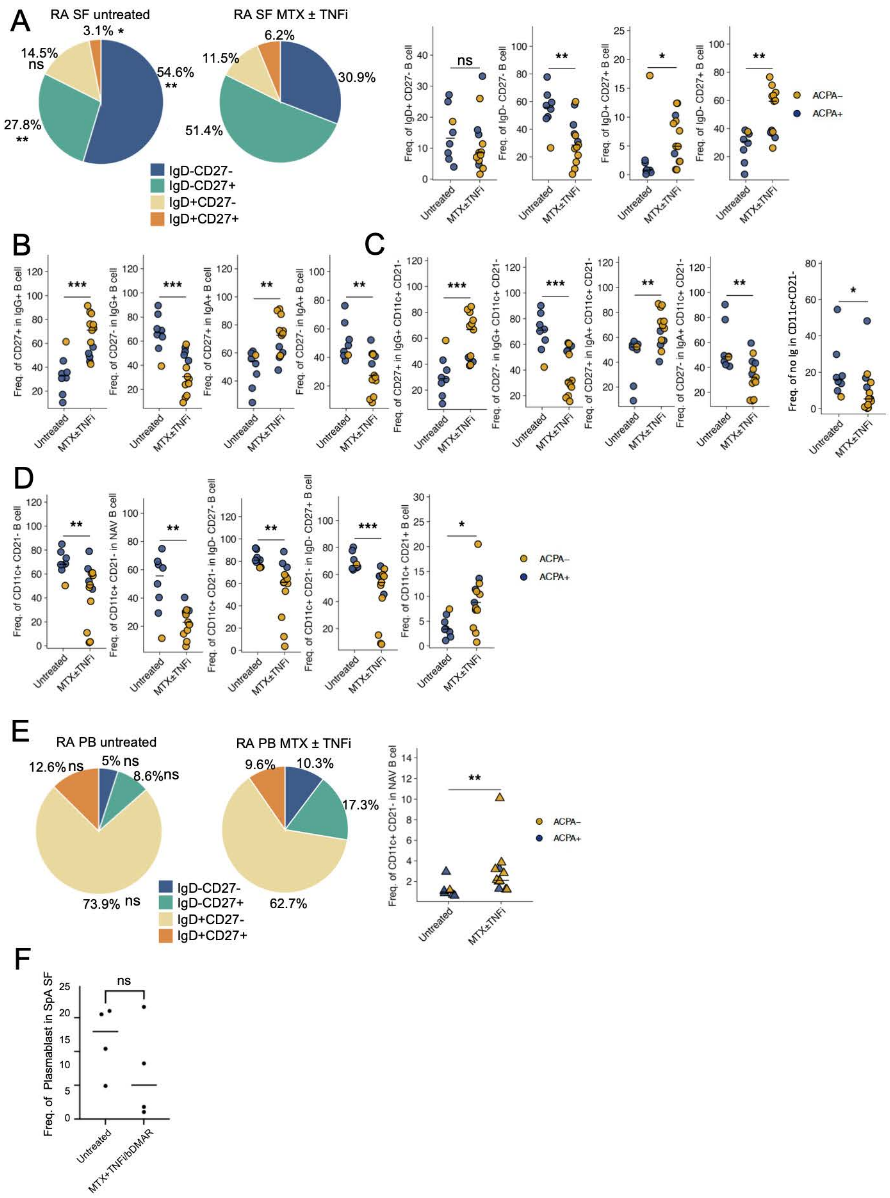


**Supplementary Fig. 10 | Effects of MTX ± TNFi treatment on B-cell subset composition in RA.**

**A**. Pie charts of the median frequencies of the four IgD/CD27-defined B-cell subsets in SF from untreated versus MTX ± TNFi-treated RA patients, with accompanying dot plots per subset (ACPA+, dark; ACPA-, orange). **B, C** Graphs comparing untreated versus MTX ± TNFi-treated RA for CD27+/CD27- within IgG+ and IgA+ B cells and for the frequency of CD11c+ CD21- B cells within the total, NAV, non-NAV, DN2 and no Ig compartments. **D**. Graphs show the proportion of CD11c+ CD21- cells within all B cells, naïve (NAV) or double negative CD27- IgD- B cells in PB from untreated versus treated RA. **E.** Pie charts show the proportion of IgD/CD27-defined subsets in PB from untreated versus MTX ± TNFi-treated RA patients; dot plot show the CD11c+CD21- in NAV B cell comparison based on treatment in RA PB. **F.** Frequency of plasmablasts in SpA SF from untreated versus MTX + TNFi/bDMARD-treated patients. Each symbol represents one patient. Mann-Whitney U test was used for statistical analysis. *P < 0.05, **P < 0.01, ***P < 0.001; ns, not significant.

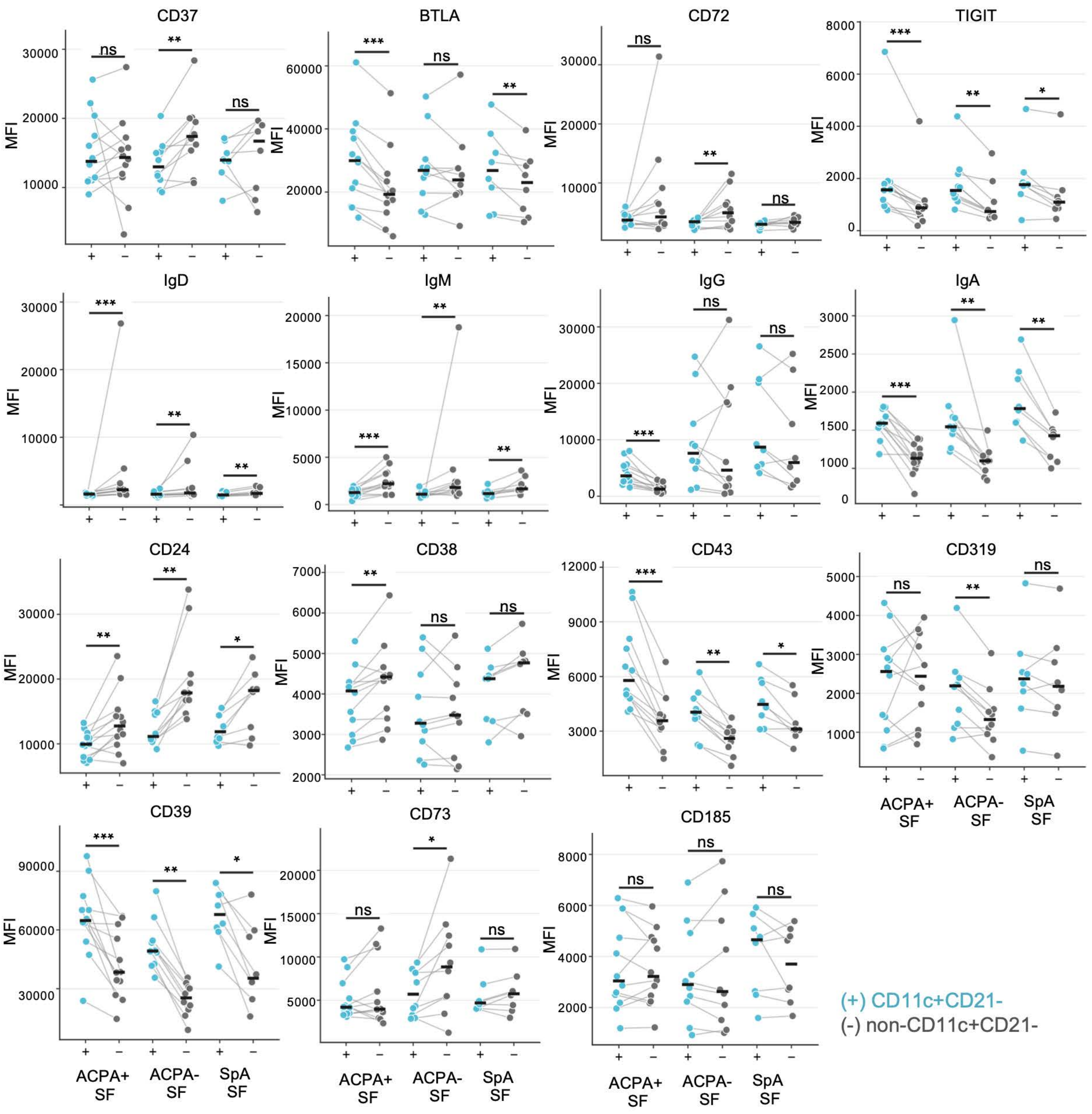


**Supplementary Fig. 11 | Extended surface-marker comparison between CD11c+CD21- and non-CD11c+CD21- B cells.**

Graphs are comparing paired MFI in CD11c+CD21- (+, cyan) versus non-CD11c+CD21- (-, grey) B cells within each SF disease group (ACPA+, ACPA-, SpA) for CD37, BTLA, CD72, TIGIT, IgD, IgM, IgG, IgA, CD24, CD38, CD43, CD319, CD39, CD73 and CD185. Paired values are connected by grey lines. Wilcoxon signed-rank test was used for statistical analysis. $^*P < 0.05$, $^{**}P < 0.01$, $^{***}P < 0.001$; ns, not significant.

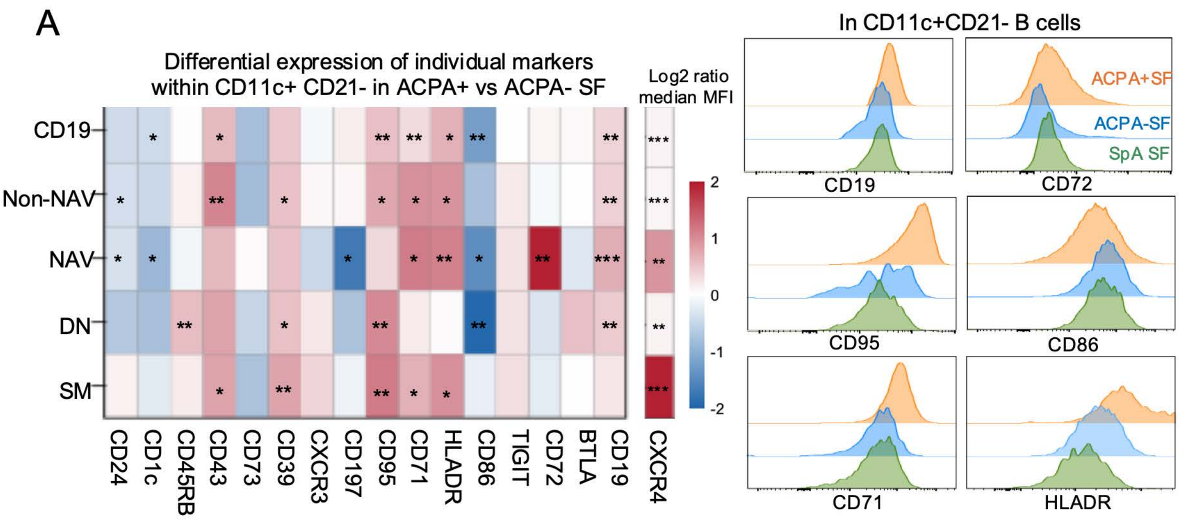


**Supplementary Fig. 12 | Differential activation-marker expression within CD11c+ CD21- B cells between ACPA+ and ACPA- SF.**

**A**. Heatmap of the $\log_2$ median-MFI ratio for individual markers within CD11c+ CD21- B cells across subsets (CD19+, non-NAV, NAV, DN, SM) from ACPA+ versus ACPA- SF. Red indicates higher expression and blue lower expression in ACPA+ RA. Representative flow cytometry histograms overlays of CD19, CD72, CD95, CD86, CD71 and HLA-DR within CD11c+ CD21- B cells across disease groups (ACPA+ SF, orange; ACPA- SF, blue; SpA SF, green) are shown at right. Wilcoxon signed-rank test was applied. *P < 0.05, **P < 0.01, ***P < 0.001.

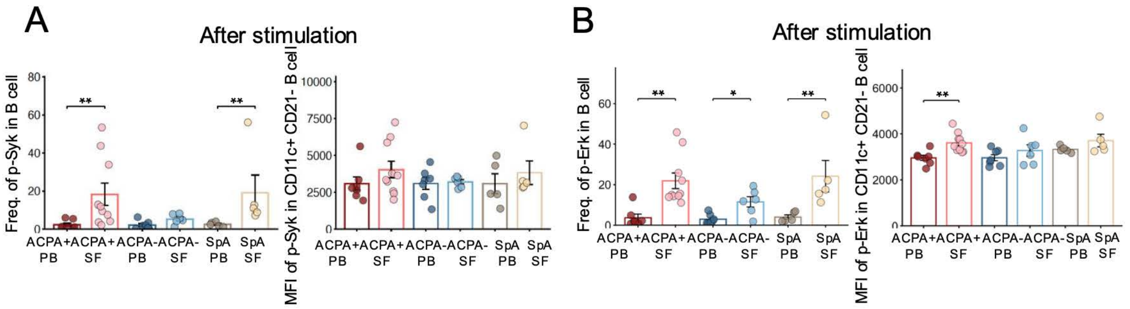


**Supplementary Fig. 13 | BCR-proximal signaling responses in B cells among ACPA+ RA, ACPA- RA and SpA**

PBMCs were stimulated with anti-Ig (Jackson Immunoresearch) for 3min and phosphorylation of Erk (p-ERK) and Syk (p-Syk) in B cells was assessed by intracellular phospho-flow. Frequency within all B cells (left) and MFI within CD11c+ CD21- B cells (right) of p-Syk (A) and p-Erk (B) after stimulation, in PB and SF across disease groups. Wilcoxon signed-rank test was applied. *$P < 0.05$, **$P < 0.01$, ***$P < 0.001$.

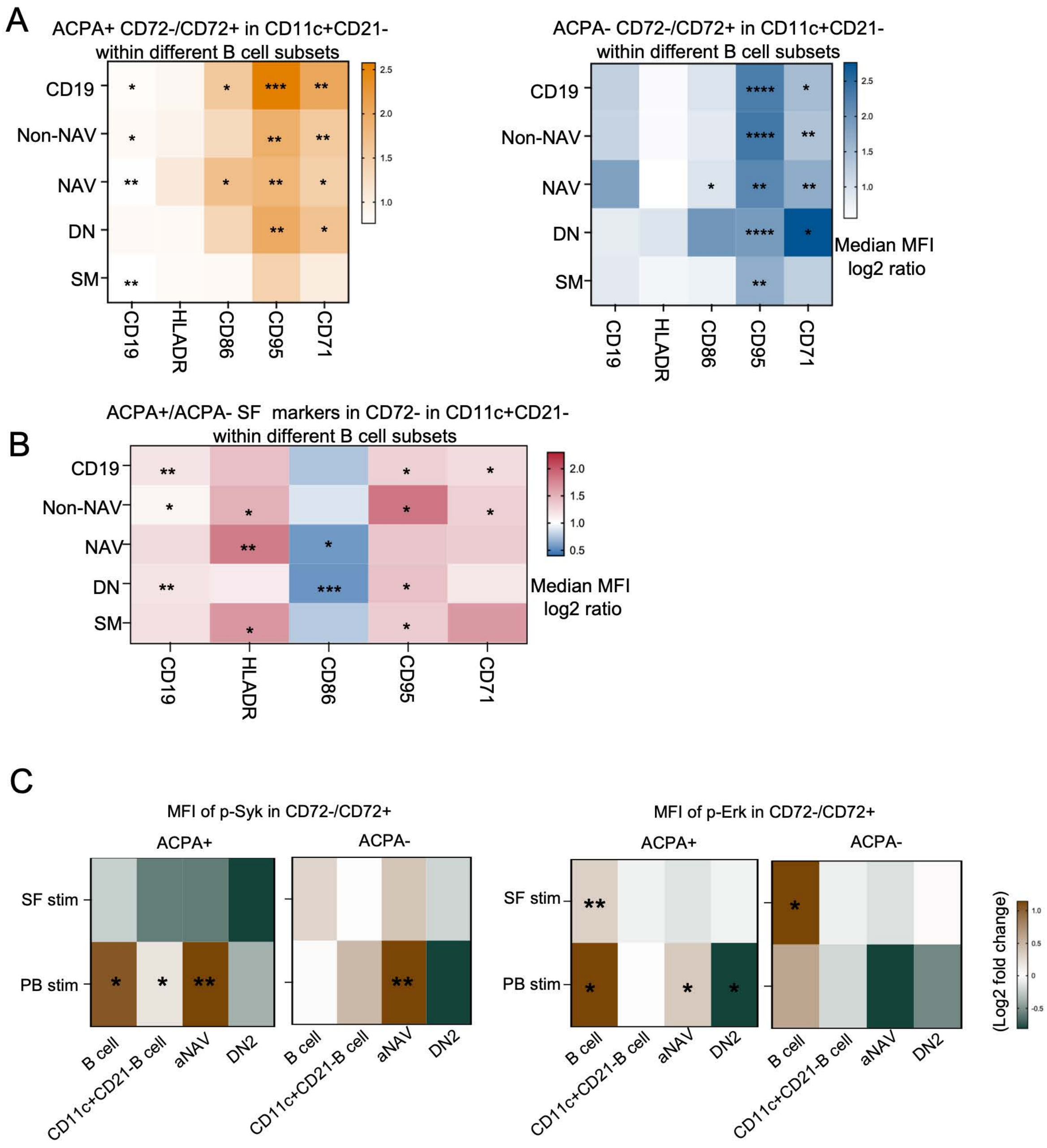


**Supplementary Fig. 14 | CD72 loss defines a broadly activated phenotype of CD11c+ CD21- B cells and elevated BCR signaling in aNAV.**

**A.** Heatmaps of the $log_2$ median-MFI ratio (CD72-/CD72+) for CD19, HLA-DR, CD86, CD95 and CD71 within CD11c+ CD21- B cells across subsets (CD19+, non-NAV, NAV, DN, SM) in ACPA+ (left) and ACPA- (right) SF. **B.** Heatmap of the ACPA+ SF/ACPA- SF $log_2$ MFI ratio for the same activation markers restricted to the CD72- subset of CD11c+CD21- B cells across subsets. **C.** Heatmaps of the $log_2$ fold change in p-Syk (left) and p-Erk (right) MFI between the CD72- and CD72+ subset within CD11c+CD21- B cells, stratified by subset (total B cell, CD11c+CD21- B cell, aNAV, DN2) and condition (SF stimulated, PB stimulated,) for ACPA+ and ACPA- samples. Wilcoxon signed-rank test was applied. *P < 0.05, **P < 0.01, ***P < 0.001.

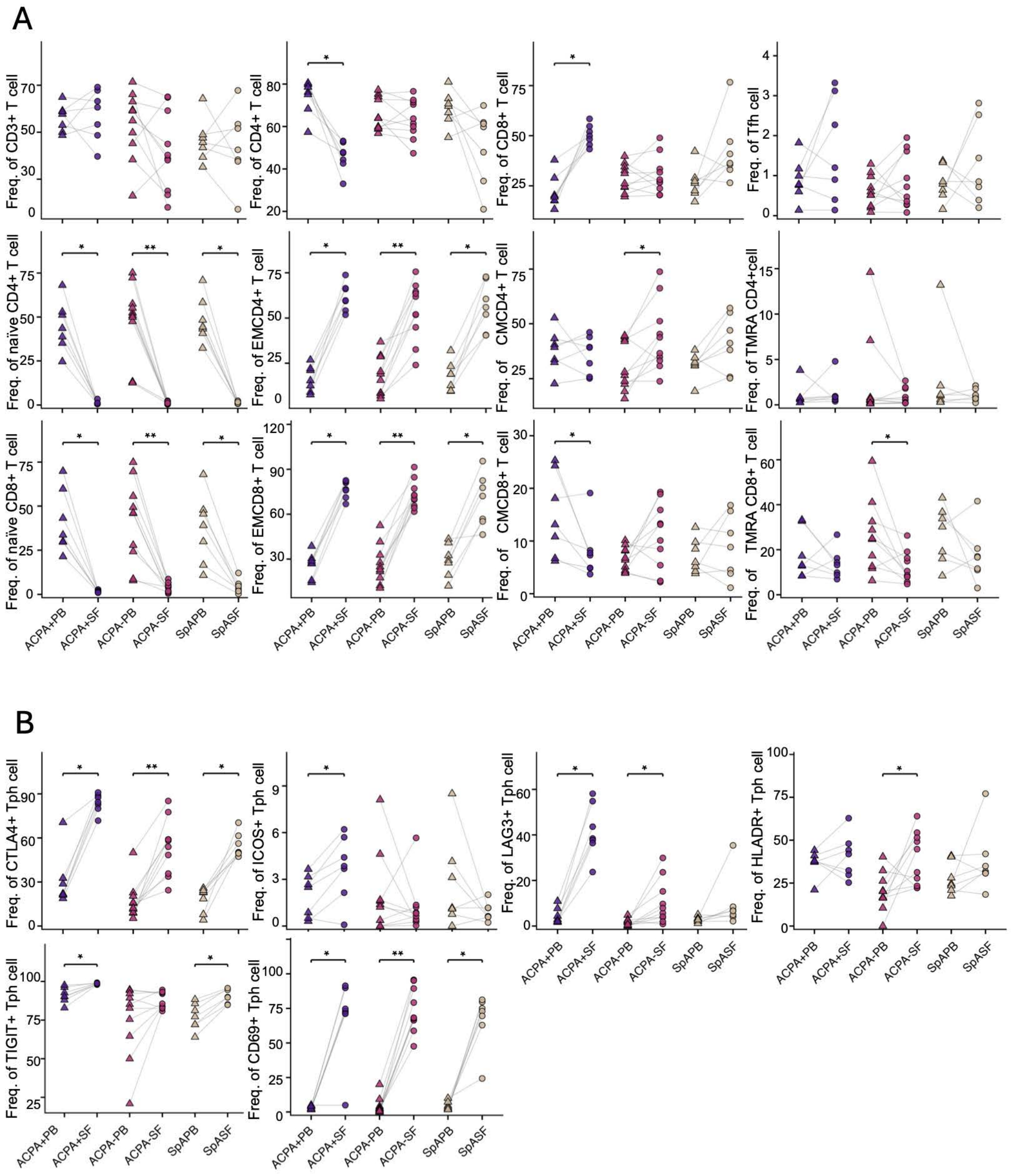


**Supplementary Fig. 15 | T-cell subset frequencies in paired blood and synovial fluid**

**A.** Paired PB-SF dot plots showing the frequency of CD3+ T cells, CD4+ T cells, CD8+ T cells and Tfh cells, and of naïve, effector-memory (EM), central-memory (CM) and TEMRA subsets within CD4+ and CD8+ T cells, across disease groups. **B**. Paired PB-SF dot plots showing the frequency of CTLA4+, ICOS+, LAG3+, HLA-DR+, TIGIT+ and CD69+ cells within Tph cells across disease groups. Each symbol represents one patient; paired samples are connected by grey lines. Wilcoxon matched-pairs signed-rank test was used for statistical analysis. *P < 0.05, **P < 0.01, ***P < 0.001.

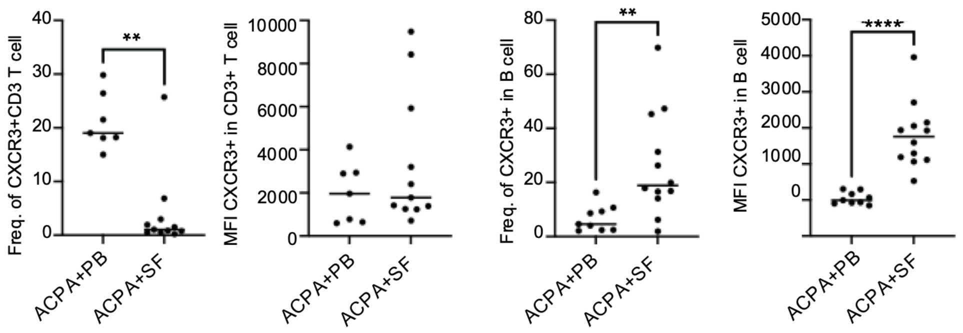


**Supplementary Fig. 16 | CXCR3 expression is decreased in synovial T cells but increased in B cells**

CXCR3 frequency and MFI comparison between PB and SF in CD3+ T cell and B cell. Line indicates median. Mann-Whitney U test was used for statistical analysis. *P < 0.05, **P < 0.01.

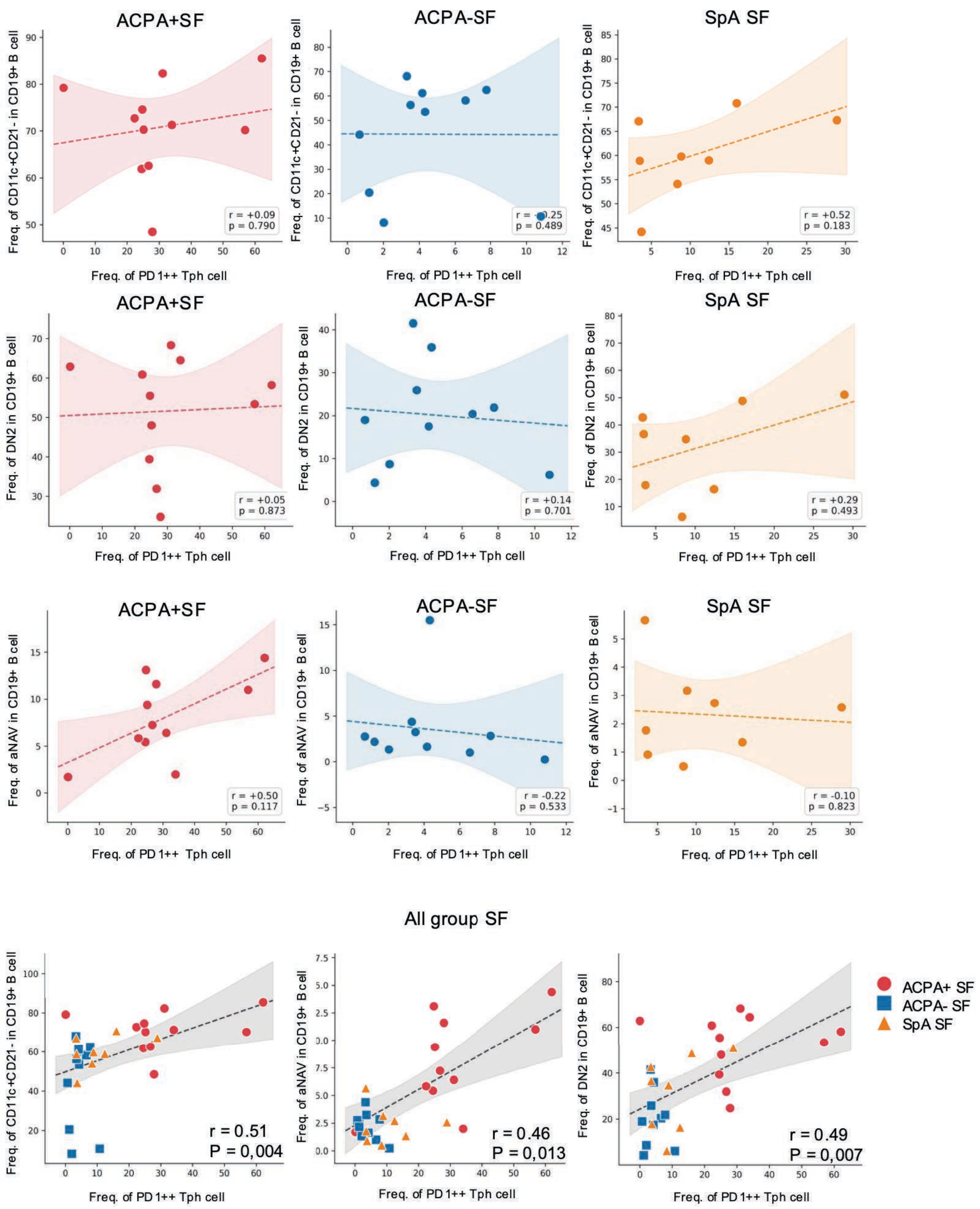


**Supplementary Fig. 17 | Correlations between PD1++ Tph cells and CD11c-associated B-cell subsets within and across disease groups.**

Correlations between the frequency of PD1++ Tph cells and the frequency of CD11c+CD21-, DN2 and aNAV cells within CD19+ B cells, shown separately for ACPA+ SF, ACPA- SF and SpA SF, and for all SF samples pooled (bottom row; ACPA+, red circles; ACPA-, blue squares; SpA, orange triangles). Shaded bands indicate the 95% confidence interval; Spearman's ρ and P values are shown.

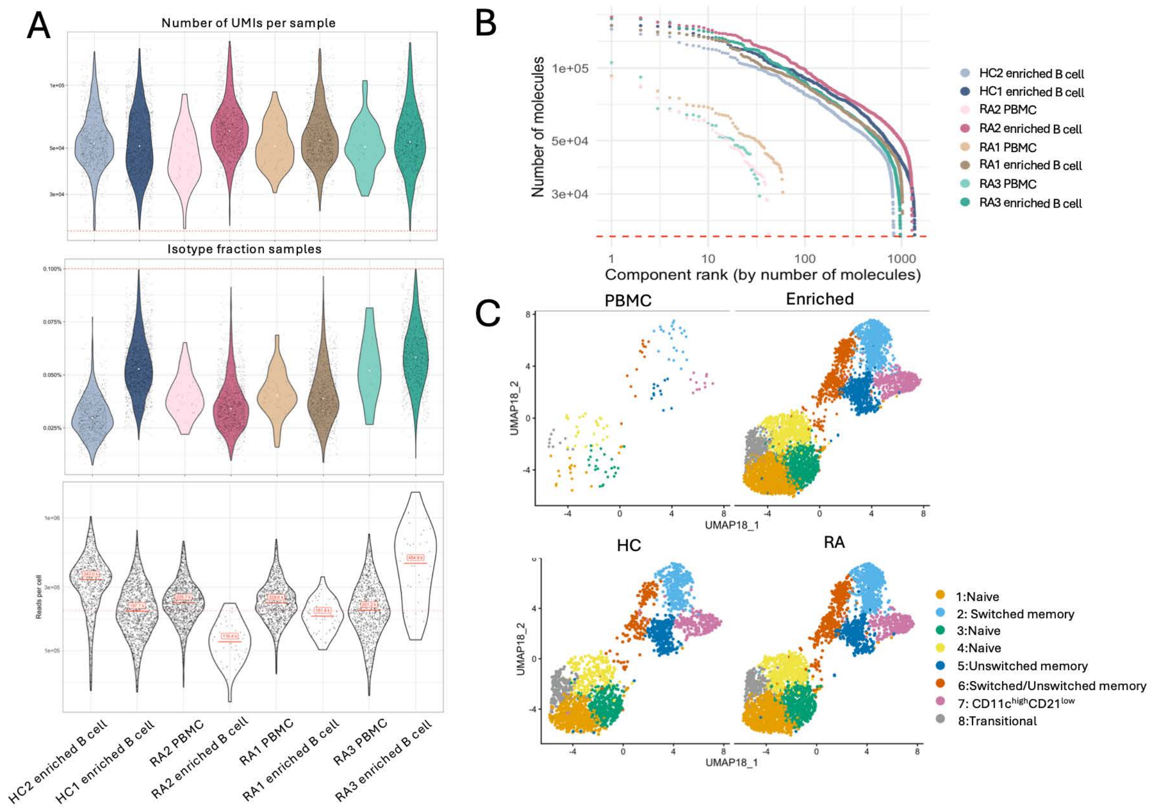


**Supplementary Fig. 18 | Quality-control metrics for PNA of B cells.**

**A.** Number of UMIs per sample, isotype fraction and reads per cell across samples. **B**. Molecule saturation (number of molecules versus component rank) per sample. **C.** UMAP embeddings of B cells split by sample type (PBMC, enriched) and by group (HC, RA), colored by cluster. Samples comprise enriched B cells and PBMCs from three untreated ACPA+ RA patients (RA1-RA3) and enriched B cells from two healthy controls (HC1, HC2).

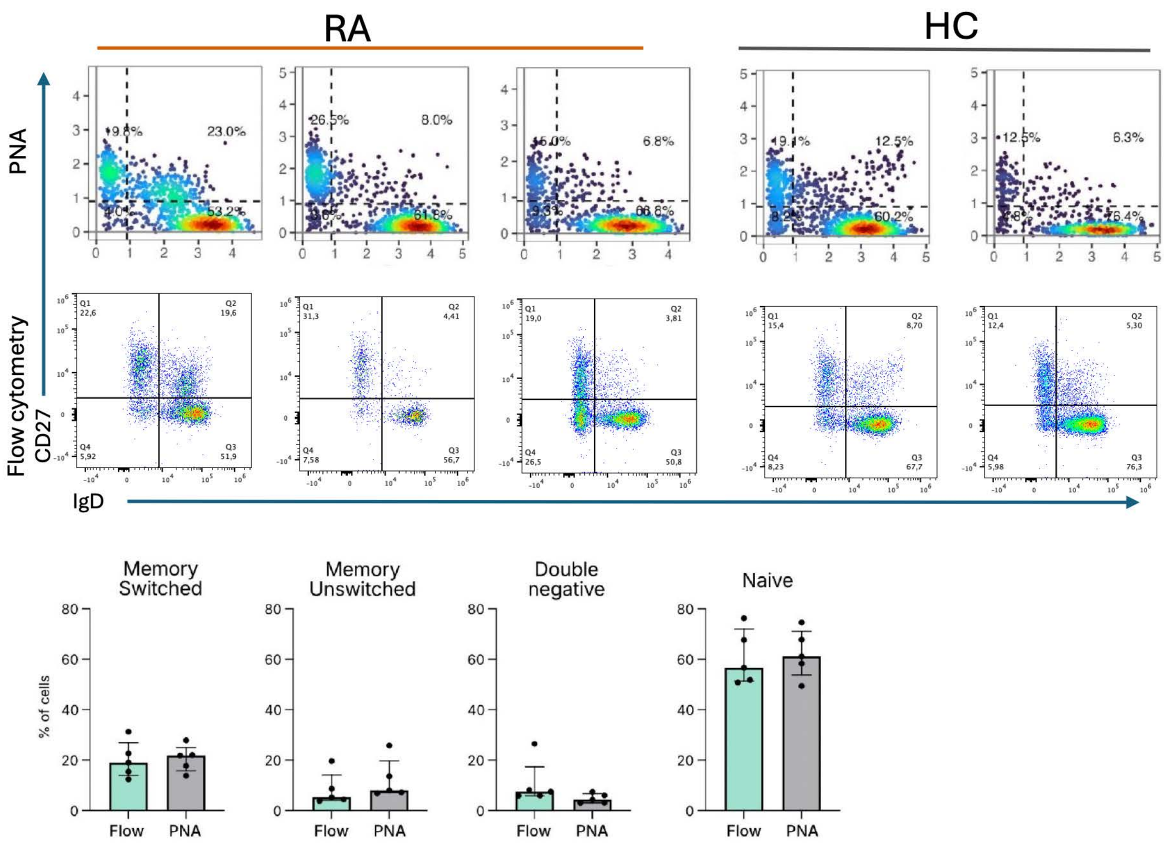


**Supplementary Fig. 19 | B-cell subset frequencies derived from the Proximity Network Assay recapitulate conventional flow-cytometry gating.**

IgD/CD27-based identification of B-cell subsets was compared between the Proximity Network Assay (PNA) and conventional spectral flow cytometry, in enriched B cells from RA patients (n = 3) and healthy controls (HC; n = 2). Top, representative IgD versus CD27 gating reconstructed from PNA surface-protein abundance; middle, matched flow-cytometry IgD versus CD27 plots for the same samples. Quadrants define switched memory (CD27+ IgD-), unswitched memory (CD27+ IgD+), naïve (CD27- IgD+) and double-negative (CD27- IgD-) B cells. Bottom, bar graphs comparing the frequency (% of cells) of switched-memory, unswitched-memory, double-negative and naïve subsets defined by flow cytometry (teal) versus PNA (grey) across all donors. Each symbol represents an individual donor; bars show mean ± s.d. Subset frequencies were concordant between the two methods, confirming that the PNA/Molecular Pixelation platform faithfully reproduces flow-defined B-cell subset composition.

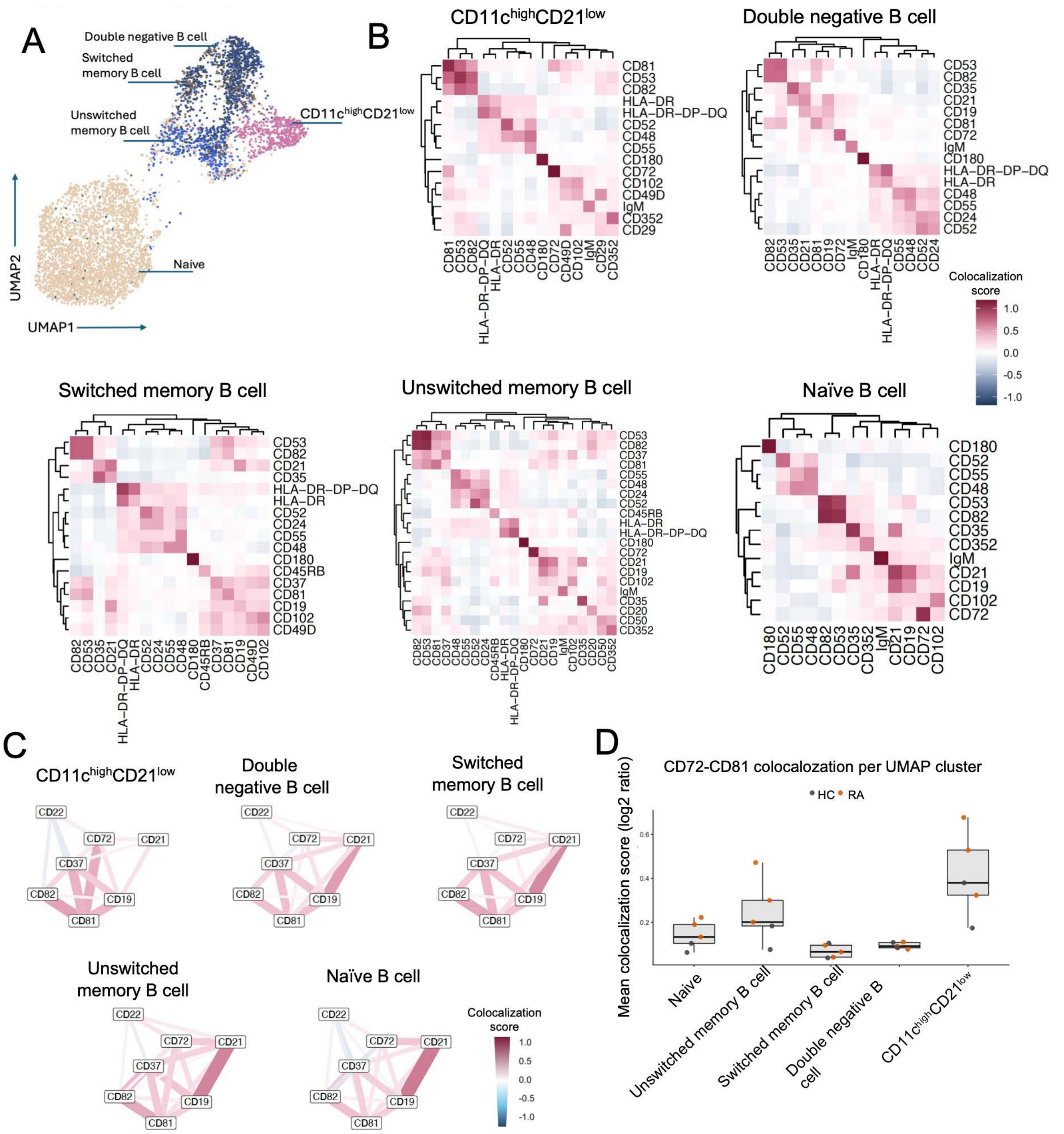


**Supplementary Fig. 20 | Proximity analysis of B cells grouped into five canonical cell types.**

**A.** UMAP of enriched B cells annotated into five cell types (naïve, switched memory, unswitched memory, double-negative and CD11chigh CD21low B cells). **B**. Clustered heatmaps of pairwise protein proximity within each cell type. (Colour represent mean $\log_2$ ratio) **C**. Network graphs of BCR-complex/tetraspanin markers (CD19, CD21, CD22, CD37, CD72, CD81, CD82) per cell type (edge colour and width represent mean $\log_2$ ratio). **D** CD72-CD81 mean co-localization score ($\log_2$ ratio) per cell type (HC, grey; RA, orange). Proximity data were pre-filtered for individual marker counts > 100 and join count > 5.

**REFERENCE**

1. Korsunsky, I., et al., *Fast, sensitive and accurate integration of single-cell data with Harmony.* Nat Methods, 2019. **16**(12): p. 1289-1296.
2. Van Gassen, S., et al., *FlowSOM: Using self-organizing maps for visualization and interpretation of cytometry data.* Cytometry A, 2015. **87**(7): p. 636-45.
